# Design of a Biohybrid Materials Circuit with Binary Decoder Functionality

**DOI:** 10.1101/2023.08.10.552766

**Authors:** Hasti Mohsenin, Hanna J. Wagner, Marcus Rosenblatt, Svenja Kemmer, Friedel Drepper, Pitter Huesgen, Jens Timmer, Wilfried Weber

## Abstract

Synthetic biology applies concepts from electrical engineering and information processing to endow cells with computational functionality. Transferring the underlying molecular components into materials and wiring them according to topologies inspired by electronic circuit boards has yielded materials systems that perform selected computational operations. However, the limited functionality of available building blocks is restricting the implementation of advanced information-processing circuits into materials. Here, we engineer a set of protease-based biohybrid modules the bioactivity of which can either be induced or inhibited. Guided by a quantitative mathematical model and following a design-build-test-learn cycle, we wire the modules according to circuit topologies inspired by electronic signal decoders, a fundamental motif in information processing. We design a 2-input/4-output binary decoder for the detection of two small molecules in a material framework that could perform regulated outputs in form of distinct protease activities. The here demonstrated smart material system is strongly modular and could be used for biomolecular information processing for example in advanced biosensing or drug delivery applications.

## 1 Introduction

A feature of every living cell is its ability to sense, process and respond to environmental stimuli.^[1]^ Engineering and rewiring the underlying molecular components using synthetic biology techniques enabled the implementation of complex information-processing circuits in living cells.^[2–9]^ Such circuits paved the way for advanced applications for example in determining and combatting multifactorial disease states^[10]^, in integrated biosensing for the detection of toxic contaminants^[11]^, in the rational design of circuits that induce cell death^[4]^, for rewiring communication between cells^[12]^, or for the design of fast biosensing logic circuits.^[13]^ In recent works, both molecular building blocks as well as engineering design concepts were transferred from synthetic biology to materials science. The functional coupling of molecular 2 sensors and actuators to polymer materials according to circuit topologies inspired from electrical engineering yielded biohybrid material circuits that perform fundamental and advanced information processing such as signal amplification using positive feedback and feed-forward loops^[14,15]^, the ability to count the number of input light pulses^[16]^, logic-based delivery of proteins^[17]^, or different logic gate designs.^[18,19]^

These material circuits have enabled applications in different areas such as the sensitive detection of drugs or toxins by signal-amplifying material systems^[14,15]^, the release of drugs on command^[20]^, or the light-controlled sequential catalysis of multi-step biochemical reactions.^[16,21]^ Moreover, these systems have found applications for instance for remodelling the extracellular matrix ECM^[22,23]^, in 3D cell-culture^[24]^, and as biosensors for the detection of environmental stimuli such as reducing agents, light and enzymes.^[25]^

Despite these advances, the design of biohybrid information-processing material circuits is limited by the scarce palette of materials-compatible biological building blocks relying mainly on mutual activation thus leaving circuit topologies excluded that require inhibitory interactions. To overcome these limitations, here we engineer a set of biomolecular switches for incorporation into biohybrid materials that can either be activated or inhibited as a function of the input signal. To demonstrate functionality of the switches and the thereby enabled novel opportunities in designing advanced information-processing biohybrid material circuits, we implement a binary 2-input/4-output decoder. Binary decoders are used to convert n-coded inputs to a maximum of 2^n^ unique outputs and are fundamental motifs in information processing. For example, binary decoders play a vital role in everyday electronic applications, moreover in medical and healthcare informatics and neuroscience studies.^[26–28]^

As functional units of these circuits we chose sequence-specific proteases, enzymes that selectively recognize their specific target peptide sequence and cleave it. Proteases have been engineered as important control elements in synthetic biology.^[29–32]^ Their activities can be tailored and engineered to respond to various inputs through fragment completion and dissociation, or substrate recruitment.^[4,33,34]^ The Potyvirus family of proteases comprises several highly specific proteases with orthogonal cleavage sites, thus allowing the use of multiple proteases within one single system.^[35,36]^ Tobacco etch virus protease (TEV) and tobacco vein mottling virus protease (TVMV) are two well-characterized examples from this family of cysteine proteases and can be engineered in their full- or split form. Splitting proteases into two parts yields inactive fragments, that however regain functionality once reconstituted by heterodimerization. Triggering such dimerization via constitutive or stimuli-responsive protein-protein interactions has resulted in the design of molecular sensors and switches to sense intracellular processes such as phosphorylation, or to act as intracellular sensor/actuator devices to control protein activity.^[37][34]^ A versatile tool to control protein-protein interactions are coiled-coil domains, short modular protein motifs that form highly specific dimers.^[4,13,38– 41]^ Another well-characterized family of proteases are caspases that play a crucial role in cell apoptosis.^[42,43]^ Caspases have been engineered and utilized as versatile tools for synthetic circuits and information-processing materials.^[14,44]^

Design-build-test-learn (DBTL) iterative workflows in synthetic biology aim to accelerate the system development and optimization by reducing laboratory cost. DBTL gains enormous productivity when supported by in-depth data analysis and mathematical modelling.^[23,45,46]^

In this work, we engineer split proteases to be inactivated by another protease via cleavage of the coiled-coil domain that interlinks the split protease fragments. We further develop material modules that release the engineered split proteases on demand in response to specific small-molecule stimuli. In the next step, we assembled the protease-based modules to fundamental logical gates emulating AND as well as NOT functionality. We finally wired these gates to a 2-input/4-output binary decoder following a DBTL approach in which the parameters to be optimized in each iteration are identified by a quantitative mathematical model.

Thanks to the engineering of inhibitory functions into protease-based signalling, novel circuit topologies as exemplified by the 2-input/4-output binary decoder become accessible for integration into materials circuits. Further, the underlying, iterative design approach guided by a quantitative mathematical model serves as blueprint for the development of different information-processing biohybrid material circuits, which enables important applications in different fields such as integrated sensors and switches, bioanalytical devices with integrated information processing, or smart drug delivery.

## 2 Results and Discussion

### 2.1. System Design and Mode of Function

In this study, we combined material modules with synthetic biological building blocks to develop biohybrid materials capable of performing fundamental information-processing operations. We designed a biohybrid binary decoder for sensing combinations of two different input molecules and converting them into 4 distinct outputs. Presence and/or absence of each input signal would result in a unique corresponding output, as shown in **Figure 1a**. For this, we chose two inputs: the antibiotic novobiocin (Novo) as input 1 (IN1) and the small-molecule ethylenediaminetetraacetic acid (EDTA) as input 2 (IN2), and four different outputs corresponding to the activities of three different proteases, or the lack thereof. The three distinct proteases comprise TEV protease, TVMV protease, and a human caspase-3 (Casp3) protease. The activity of proteases is monitored by measuring the time-resolved changes in the concentration of their respective cleaved substrates. Relative enzyme units report the amount of substrate cleaved per min. The overall circuit design of the 2:4 binary decoder is shown in Figure 1b. In the absence of both input molecules, the output if OFF. In the presence of only novobiocin, TEV protease is released. Similarly, EDTA as sole input triggers the release of TVMV only. When both inputs are present, both TEV and TVMV proteases are released and eventually trigger release of Casp3. However, upon release of Casp3, the proteases TEV and TVMV are cleaved and thus inhibited, leaving Casp3 as the main remaining output.

**Figure 1.**
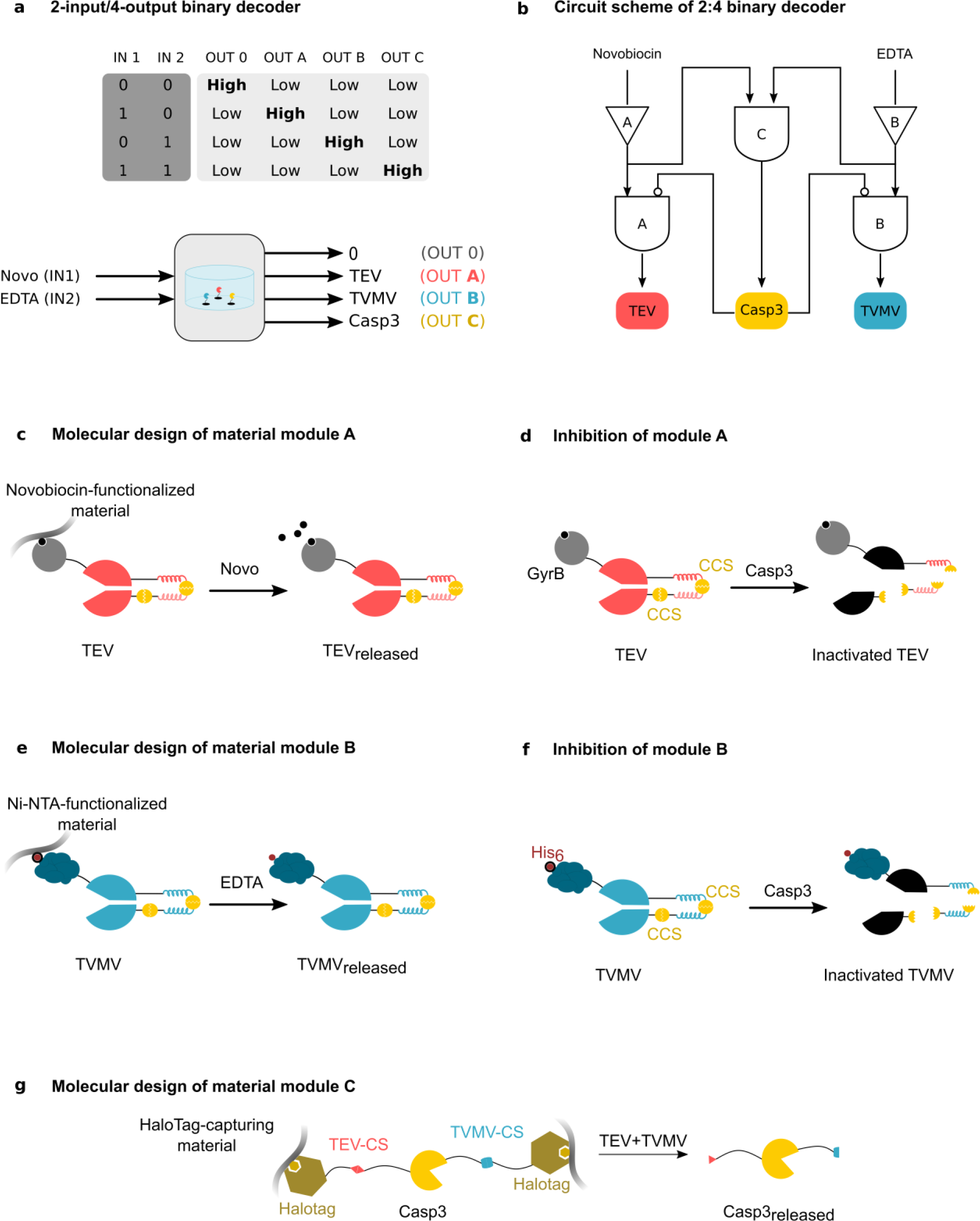
Design of a binary decoder. a) Principle of a 2:4 binary decoder with two input (IN) and four output (OUT) signals and the different input/output combinations (upper panel). The addition of novobiocin (Novo, IN1) and EDTA (IN2) serve as input. The four different output states are represented by the release of distinct proteases, none (OUT 0), TEV protease (OUT A), TVMV protease (OUT B), or Casp3 protease (OUT C). The different proteases are introduced in the material modules A, B, C (lower panel). b) Circuit scheme of the 2:4 binary decoder. Addition of novobiocin releases TEV protease (module A). Addition of EDTA results in TVMV release (module B). Only when both novobiocin and EDTA are present, released TEV and TVMV release Casp3 from material module C. Subsequently, released Casp3 module inactivates TEV and TVMV signals, resulting in the Casp3 signal as the main output. c) Molecular design of material module A. Module A consists of a split-TEV protease, fused to GyrB. GyrB serves as anchor for coupling to the polymer support, novobiocin-functionalized-magnetic crosslinked agarose. Upon addition of free novobiocin, split TEV is released. d) Inhibition of module A. Casp3 protease cleaves its cleavage site (CCS) in the coiled-coil regions of the construct and thereby inhibits TEV activity by dissociating its split components. e) Molecular design of material module B. His_6_-tagged split-TVMV is bound to Ni^2+^ -NTA-functionalized crosslinked magnetic agarose via its His_6_-Tag and is released from the material in the presence of Ni^2+^ -chelating EDTA. f) Inhibition of module B. Casp3 inactivates TVMV protease by cleaving its cleavage site in the linkers connecting the TVMV split parts, causing the split parts to dissociate. g) Molecular design of material module C. Material module C is a Casp3 protease genetically fused to N-/ and C-terminal HaloTags and covalently bound to its material support, HaloTag-capturing magnetic particles encapsulated with microporous cellulose. Only when both TEV and TVMV are present, the linkers between Casp3 and HaloTags containing TEV-CS and TVMV-CS are cleaved, respectively, to release Casp3.

This design requires building blocks that can either be released/activated or inhibited in response to a distinct stimulus. We implemented this dual-control into the corresponding modules as follows: Module A is composed of a TEV protease genetically fused to bacterial subunit gyrase B (GyrB) to enable its binding onto novobiocin-functionalized magnetic crosslinked agarose with high affinity (K_d_ value 1-2 ×f10^-8^ M^[47]^). This module releases TEV protease in the presence of free novobiocin (Figure 1c). TEV protease is designed in its split-form and the N-/ and C-terminal parts are joined together via coiled-coil regions AP4 and P3 introduced by Fink et al..^[13]^ The linker between the split regions of TEV contains two Casp3-cleavage sites (CCS), one between the two coil regions and the other one in the linker fusing AP4 to the C-terminal of TEV. TEV is expected to be inhibited by the addition of Casp3 protease by cleaving CCS to dissociate and inactivate the split TEV parts from each other (Figure 1d).

Module B consisted of TVMV protease, where we tested its immobilization and release via two different mechanisms, either via chelation mechanism, or via addition a metabolite-responsive allosterically regulated DNA-binding protein (Figure S2). The first mechanism comprised TVMV protease coupled through its His6-Tag onto magnetic crosslinked agarose functionalized with Ni^2+^ -NTA with an affinity of 1-2 ×f10^-8^ M.^[48]^ Upon addition of EDTA, Ni^2+^ is chelated and TVMV is released (Figure 1e, Figure S2a). The second mechanism included TVMV protease, genetically fused to a single-chain uricase oxidase repressor protein (scHucR).^[49,50]^ HucR binds to its cognate operator DNA-sequence, hucO. The HucR-hucO complex binds via biotin functionalization of hucO onto streptavidin-functionalized magnetic macroporous cellulose material. By adding uric acid, HucR undergoes a conformational change and the HucR-TVMV fusion is released from its material support (Figure S2b). Due to the better performance of the chelation mechanism, and higher stability of His_6_-Tag compared to scHucR upon longer storage times (Figure S10), the chelation-based release mechanism was chosen for subsequent experiments. The engineered split TVMV is similar to split TEV, with the difference that the split protease is TVMV and the coiled-coil regions are an orthogonal pair to the ones used for TEV, called P9mS and A10.^[13]^ Inhibition is induced by Casp3 cleaving the CCS in the linker and therefore enables split TVMV dissociation and inactivation (Figure 1f). For module C, we genetically fused Casp3 to HaloTags on both N-/ and C-termini, via linkers containing cleavage sites for TVMV and TEV proteases, TVMV-CS and TEV-CS, respectively. The HaloTags allow a covalent coupling to the material support, HaloTag-capturing magnetic particles encapsulated with microporous cellulose. In this configuration, Casp3 is released only in the presence of both TEV AND TVMV (Figure 1g).

### 2.2. Synthesis and Characterization of each Material Module

Prior to decoder assembly, we evaluated the release of proteases from each module individually. Each protein building block was coupled onto crosslinked magnetic agarose and subsequently immobilized in a multiwell plate via a magnet placed under the plate. The response of each module was monitored upon being exposed to the different input combinations. Modul A comprised magnetic crosslinked agarose that was functionalized with novobiocin. Novobiocin enabled the coupling of TEV construct via its GyrB sequence onto the solid support. The functionality of this module was evaluated by adding novobiocin or EDTA to the system and measuring the output TEV. The release of TEV only in the presence of novobiocin indicated specificity to this input (**Figure 2a**, see **Figure S3a** for the time-course of TEV release). Similarly, module B consisted of split-TVMV containing a His_6_-Tag, by which this module was coupled onto Ni^2+^ -NTA-functionalized magnetic crosslinked agarose. Module B as well showed a response in form of TVMV activity, only in the presence of EDTA, without showing non-desired TVMV activities in presence of novobiocin or without any inputs (Figure 2b, see Figure S3b for the time-course of TVMV release). In module C, Casp3 was N-/ and C-terminally coupled to HaloTag-capturing magnetic crosslinked agarose. The release of Casp3 only in the presence of both TEV and TVMV demonstrates the AND-type functionality of this configuration (Figure 2c).

**Figure 2.**
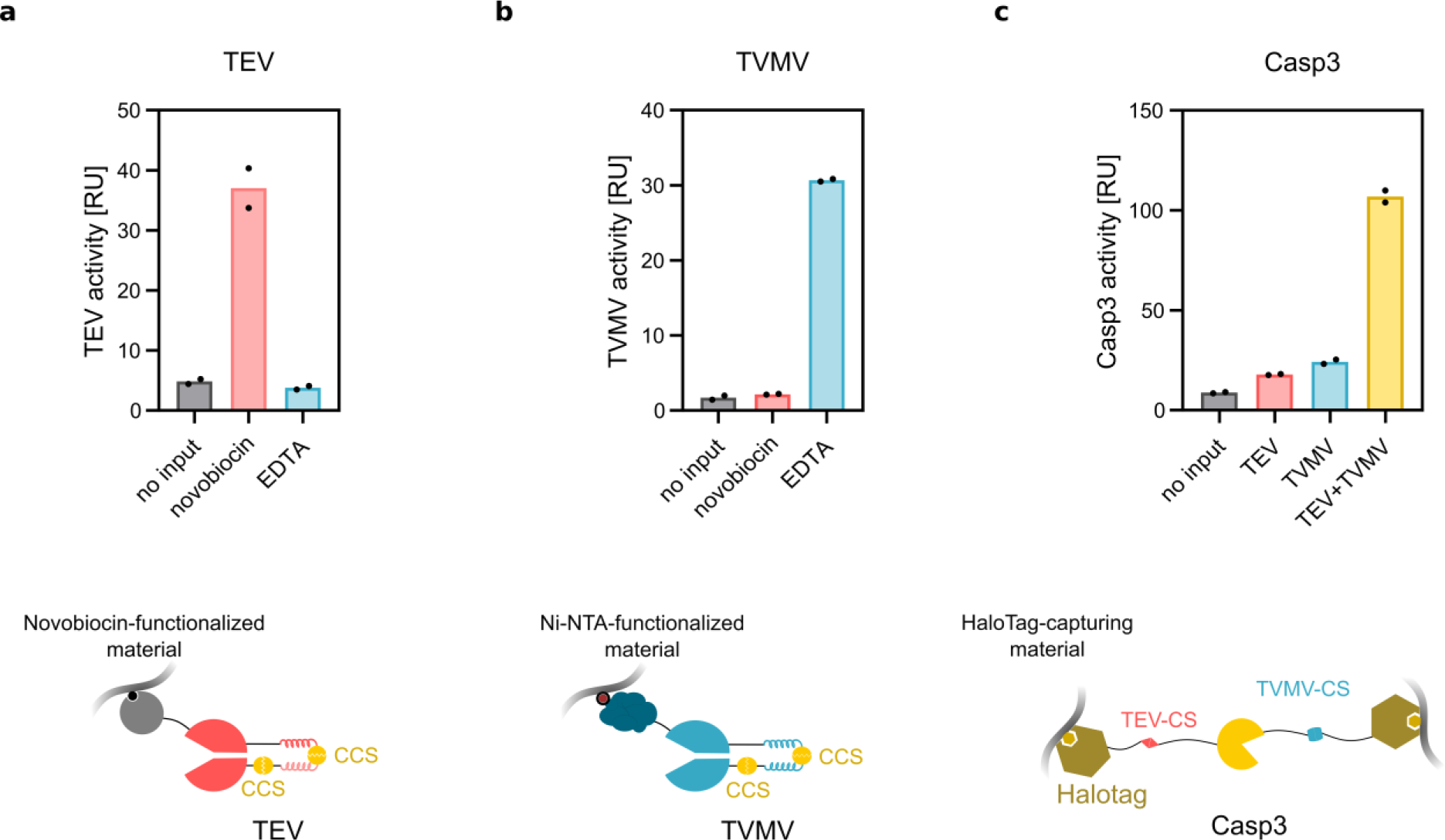
Development and characterization of the individual protease-based modules. a) Release of split TEV protease from module A in response to different inputs. 60 µg of agarose-immobilized split TEV protease was incubated in 550 µL assay buffer in the presence or absence of novobiocin (53.6 µM) or EDTA (10 mM) for 2 h prior to scoring TEV activity in the supernatant. b) Release of split TVMV protease from module B in response to different inputs. 30 µg of agarose-immobilized split TVMV was incubated in 550 µL assay buffer in the presence or absence of novobiocin (53.6 µM) or EDTA (10 mM) for 2 h prior to scoring TVMV activity in the supernatant c) Release of Casp3 from module C in response to different inputs. 40 µg of agarose-immobilized Casp3 was incubated in 550 µL assay buffer in the presence or absence of TEV (60 µg), TVMV (30 µg) or both for 2 h prior to scoring Casp3 activity in the supernatant. Bar charts indicate the mean protease activities of two separate biological replicates.

Once the functionality of all three modules was validated, the decoder system was assembled. To obtain quantitative insight into the complex interplay of all modules, a mathematical model was developed and parameterized based on the characterization of the individual modules. Detailed description of the generated mathematical model can be found in the Supplementary Information. The development of the system followed iterative DBTL cycles where in each step the mathematical model was used to quantitatively characterize the system’s performance and guide the next iteration (**Figure S1**).

### 2.3. Binary Decoder One-pot System Assembly

To generate a system with 3 distinct material modules that can interact with each other for signal processing and material-material communication, we designed an integrated reaction system. Here, the presence of all modules in a mixture of material-bound protein building blocks would lead to non-desired interactions since proteins immobilized onto materials could already interact with each other when the materials get in close contact. To enable material-material communication while physically separating the material modules, we designed a plate stand suitable for a 12-well plate, in which we incorporated three bar magnets below each well (**Figure 3a**). Hereby, the different modules can be immobilized on one magnet each thus preventing material-material contacts while allowing communication via released protein components. For the whole decoder system set-up, we first prepared each module individually, and subsequently assembled all three modules each on top of one magnet bar each in a single reaction well. We studied the decoder response at four different conditions: no input, novobiocin (53.6 µM concentration), EDTA (10 mM concentration), or both novobiocin and EDTA and measured all three output signals (TEV, TVMV, and Casp3 protease activity) for each condition over time. The kinetic data was used to parametrize a quantitative mathematical model (**Figure S4**) that is based on ordinary differential equations (ODEs), see Supplementary Information for a detailed model description. The released enzyme activities after 5 h upon reaching a plateau for TEV and TVMV are shown in Figure 3b (see Figure S4 for kinetic data) as a function of the combination of the novobiocin / EDTA input. In the condition of no input, the output signals for all three proteases were at background level, representing the off-state. In the presence of novobiocin, TEV output was triggered whereas addition of EDTA resulted in the release of TVMV. The simultaneous release of TEV and TVMV by the addition of novobiocin and EDTA further triggered the release of Casp3. The released Casp3 was expected to inhibit and lower the activities of the other two proteases by cleaving the CCS in the linkers connecting the split parts of TEV and TVMV proteases. However, neither TEV nor TVMV activity declined in this configuration compared to the novobiocin- / EDTA-only conditions. We thus set out to perform a second iteration of the design-build-test-learn cycle.

**Figure 3.**
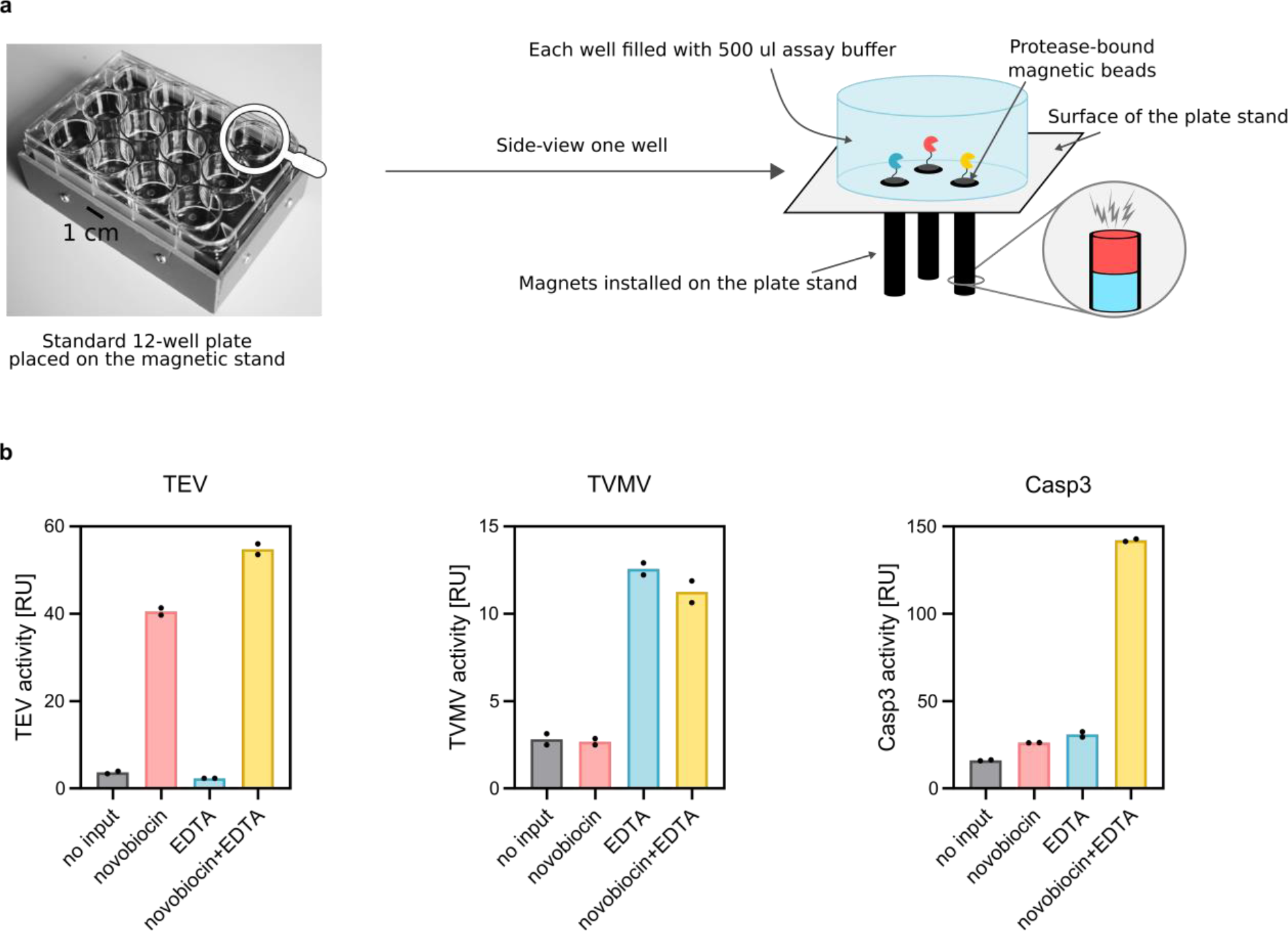
Decoder system assembly. a) Design of the magnet-based system for separating the different modules. Magnets (4 mm in diameter) are fixed and installed on the plate holder, and for each experiment a 12-well plate is placed above the magnetic stand. Magnetic beads, each functionalized with their respective proteases, Modules A, B, and C, were placed above a single magnet each in a single reaction pot, and thereby physically separated from one another while allowing communication between the modules via released proteases. b) Response characteristics of the decoder circuit. For each condition, 60 µg of agarose-immobilized split TEV, 30 µg of agarose-immobilized split TVMV, and 40 µg of agarose-immobilized Casp3 were immobilized on their respective magnets in 550 µL assay buffer. The four conditions comprise: no input, novobiocin (53.6 µM), EDTA (10 mM), or novobiocin (53.6 µM) + EDTA (10 mM). Samples were taken from the supernatant for each condition, and all three protease activities were measured for each input. Data indicate the protease activities after 5 h, bar charts show the mean of two independent biological replicates for each condition.

### 2.4. Optimizing Casp3-based inactivation of TEV and TVMV

The data in Figure 3 did not show a decrease in TEV and TVMV activity for the novobiocin+EDTA conditions despite the presence of released Casp3 in the supernatant. For TEV-containing module A, we hypothesized that an insufficient separation of the split TEV subunits could impede inactivation. To promote separation and to support inhibition, we engineered a displacer module consisting of an inactive C-terminal portion of TEV protease fused to a coiled-coil domain (P4)^[13]^ with a higher affinity to the coiled-coil counterpart fused 11 to the TEV N-terminus (**Figure 4a**). To test whether addition of the displacer supported TEV inactivation in the presence of Casp3, we compared the activity of the intact TEV variant and Casp3-treated TEV in the presence or absence of the displacer. Indeed, the displacer module significantly reduced the activity of Casp3-cleaved TEV compared to the intact variant (Figure 4a).

**Figure 4.**
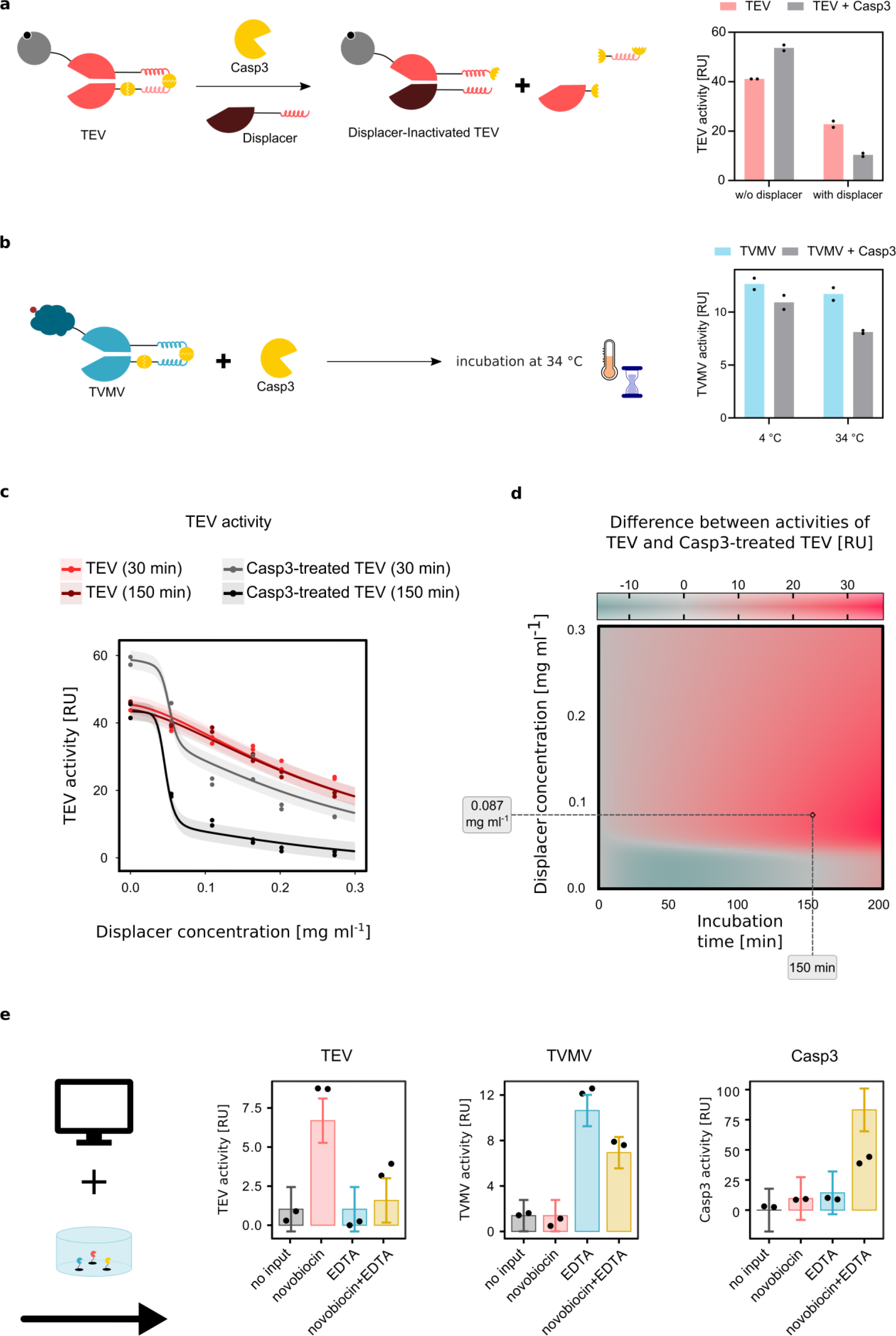
Model-guided optimization of the decoder response. a) Displacer-supported Casp3-based TEV inactivation. Mechanism of action of the displacer module (left panel). 30 µg of 14 TEV was incubated in the presence or absence of 30 µg Casp3 in 300 µL assay buffer for 30 min at 34 °C in the presence or absence of 60 µg displacer prior to analysing TEV activity. b) Casp3-based TVMV inactivation. 30 µg of TVMV was incubated in the presence or absence of 3 µg of Casp3 in 300 µL assay buffer at either 4 °C or 34 °C for 1 h prior to analysis of TVMV activity. c) Dose-response analysis of the effect of the displacer on TEV activity. 30 µg of TEV was incubated in 300 µL assay buffer in the presence or absence of 3 µg Casp3 together with the indicated displacer concentrations for 30 min or 150 min prior to quantifying TEV activity. In order to be coherent with the complete decoder setup, the buffer further contained 53.6 µM novobiocin, and the buffer for Casp3-treated conditions was further supplemented with 15 µg TVMV, 53.6 µM novobiocin, and 10 mM EDTA. Experimental data points, the mathematical model fit, and respective error bands are shown. d) Optimization of displacer treatment. The difference between the activities of TEV and Casp3-treated TEV (RU_TEV_ − RU_TEV+Casp3_) was calculated using the mathematical model as a function of incubation time and displacer concentration. The resulting value is represented by the colour in the heatmap. The black rectangle shows the chosen condition for the subsequent experiment. e) Functionality of the optimized 2-input/4-output binary decoder. The decoder was assembled by immobilizing modules A, B, C containing 60 µg TEV, 30 µg TVMV, and 40 µg Casp3, respectively, above one magnet each in 550 µL assay buffer. The samples were incubated for 300 min in the presence or absence of 53.6 µM novobiocin or 10 mM EDTA or both and subjected to incubation at 34 °C with displacer (0.087 mg mL^-1^) for 150 min prior to quantifying TEV, TVMV and Casp3 activity. The bars represent the predictions of the mathematical model together with the predicted error bars, the experimental values are indicated by the black dots.

For TVMV we have observed a decrease in activity in the complete assembly in the presence of EDTA (Figure S4) which was not observed when TVMV was released from module B in the absence of the other modules (Figure S1b). We thus hypothesized that released TVMV might partly be cleaved and de-activated by Casp3 immobilized in module C. Therefore, the dynamic range between the activities of TVMV cleaved by immobilized-Casp3 and TVMV cleaved by released-Casp3 needs to be increased. To increase TVMV inhibition upon release of Casp3 (induced by addition of novobiocin and EDTA) in relation to cleavage by immobilized Casp3 only (induced by addition of EDTA only), an additional incubation step could be introduced, where the supernatant would be removed from the material module-containing plate and incubated separately. We tested the possibility of an additional step by incubating TVMV in the presence or absence of Casp3 at either 4 °C or 34 °C for 1 h prior to scoring TVMV activity (Figure 4b). The ratio of TVMV activity in the absence or presence of soluble Casp3 was strongly increased by this additional incubation at 34 °C.

With these findings, for the next cycle of decoder development, we added an additional incubation step of all samples outside the material-module containing plate at 34 °C in the presence of 0.054 mg mL^-1^ displacer for 30 min (Figure S5a). When running the decoder experiment, TEV protease activity was at its highest in the presence of novobiocin, and a decrease in activity was observed in the presence of novobiocin and EDTA. Similarly, TVMV activity was at its highest in the presence of EDTA and decreased in the presence of both novobiocin and EDTA. For Casp3, the highest activity was seen upon addition of both inducers, novobiocin and EDTA. The time-resolved activity data of all output proteases are shown in Figure S5b. The figure further shows the fits of the mathematical model to the experimental data which were used to parameterize the model. The observed response characteristics to the input parameters reflect the functionality of the 2-input/4-output binary decoder. However, to further increase the difference between the high- and low-state signals, we entered an additional iteration of the DBTL cycle.

To this aim, we first evaluated the influence of the displacer module as a function of concentration and incubation time in detail in a pre-experiment. For this, TEV and Casp3-treated TEV were incubated with different amounts of the displacer module for 30 or 150 min (Figure 4c). The activity of cleaved TEV decreased strongly with increasing displacer concentrations. Also, for uncleaved TEV a decrease was observed albeit only at higher displacer concentrations. The same experiment was carried out for module B, but no statistically significant dependence on displacer amounts was observed for TVMV (Figure S6). To identify optimal conditions for the incubation step, we parameterized the mathematical model using the data from the displacer dose-response curves from Figure 4c. We used the model to calculate the difference between the activities of TEV and Casp3-treated TEV as a function of incubation time and displacer amounts (Figure 4d). We experimentally evaluated the prediction by choosing the following conditions for the additional incubation step in the overall decoder experiment: displacer concentration of 0.084 mg mL^-1^ and an incubation time of 150 min (indicated by a black rectangle in the heatmap in Figure 4d). These conditions are expected to reach a clear difference in activity between uncleaved and cleaved TEV while avoiding too long post sample incubations. We aimed to test this condition in the whole decoder system. Prior to the validation experiment, we predicted the decoder response at this condition by the mathematical model, the predicted values are shown in Figure 4e. This condition was subsequently evaluated experimentally (data points in Figure 4e). The time-resolved prediction and experimental data are presented in Figure S7. The observed activities agree with the model predictions and correspond to the anticipated response of the 2-input/4-output binary decoder with TEV, TVMV or Casp3 being at the high state in the presence of novobiocin, EDTA, or novobiocin and EDTA, respectively.

## 3 Conclusions

In this study, we engineered proteases into biohybrid material frameworks, and equipped them with sensory, regulatory, and communicational functions towards a 2-input/4-output binary decoder circuit design. Key for the development of the decoder material system was the development of in-vitro functional molecular switches that allow both inducible release and inducible de-activation. This combined positive and negative regulation performance opens the door to the implementation of versatile information-processing functionalities in materials systems. In the development of functional network topologies, mathematical model-guided DBTL-cycles are an efficient approach. The model parameterized with the experimental data of iteration i can be used to predict the system’s performance in iteration i+1 thus allowing the in-silico sampling of the design parameter space and the efficient prediction of parameters that contribute to desired network functionality. This work benefited from the detailed characterization by the mathematical modelling that first guided the optimization of DBTL iteration cycles towards addition of an incubation step to the workflow, and subsequently by generating a prediction heatmap, directed the DBTL iterations towards conditions resulting in a decoder functionality.

The approach shown here offers a versatile platform for the generation of biocomputing materials systems, where all the building blocks can be chosen from the library of synthetic biological building blocks or be engineered towards the desired functionalities. For example, the input molecules can freely be chosen from a toolbox of protein-interacting molecules. The latter element can on the one hand couple the biomolecule on solid supports, and on the other hand enable its stimuli-responsive release. The release can be based on various mechanisms: i) competition, for instance drug-target interactions or antibody-antigen interactions by the addition of free drug or antigen.^[51,52]^ ii) allosteric regulation of protein conformation, for instance repressor proteins in response to their inducers^[50,53]^, or protein-protein interactions based on optogenetic switches responsive to different wavelengths.^[16,54–56]^ iii) chelation of metal ion complexes as shown in this study. vi) Hydrolysis, for example peptide cleavage by protease enzymes^[15]^, or nucleic acid cleavage by CRISPR-Cas systems.^[57,58]^ Moreover, the on/off switch element can be chosen from a huge library of proteases. Apart from TEV and TVMV, split versions of other plant virus proteases have as well been described, such as turnip mosaic virus (TUMV), sunflower mild mosaic virus (SuMMV), southern bean mosaic virus (SbMV), or plum pox virus (PPV) proteases.^[13,59]^ These proteases and other split proteases offer a toolbox of orthogonal on/off switches that can be integrated into advanced materials systems performing biomolecular computations. Hence, the fundamental concept of this work can be expanded to more complex circuits benefiting from the free choice of all three main elements: input molecules, recognition and immobilization element, and finally orthogonal and flexible output proteases of choice.

Unlike stimuli-responsive materials, such biocomputing materials systems exhibit programmability.^[60]^ Although the implementation of such systems within a patient might be challenging, they may find various applications in therapies or as intelligent sensors where a specific combination of biomarkers is indicative of a disease. For example, simultaneous monitoring of elevated levels of both glucose and ketone bodies is extremely important in diagnosis and treatment of diabetic ketosis.^[61,62]^ Simultaneous detection of specific biomarkers, where a combination of them is indicative of a tumour for example, could release the desired anti-tumour compounds on demand.^[18]^

The concept introduced here highlights materials systems that are capable of receiving environmental cues of choice, processing information via communicating with each other, and producing regulated outputs. This concept lays the foundation for the development of diverse information-processing biohybrid materials that pave their way towards analytical and engineering sectors for intelligent, information-integrating and -processing multi-input, multi-output diagnostic and sensory biomaterial systems.

## 4. Experimental section

### Cloning of the plasmids

The amino acid sequence of all plasmids and their description are listed in Supplementary Information **Table S1**. Constructs were cloned by Gibson assembly^[63]^.

### Protein production and purification

All recombinant proteins were produced in *Escherichia coli* BL21(DE3) pLysS (Invitrogen) transformed with respective plasmids. Bacteria were grown at 37 °C and 150 rpm in 1 L shake flasks containing LB medium supplemented with 36 µg μL^−1^ chloramphenicol and 100 µg μL^−1^ ampicillin. Once OD_600_ reached 0.6, induction was started by addition of 1 mM IPTG and incubation was done under different conditions as described in **Table S2**. Cells were harvested by centrifugation at 6000 ×f*g* for 10 min, resuspended in 35 mL Ni Lysis buffer (50 mM NaH_2_PO_4_, 300 mM NaCl, and 10 mM imidazole, pH 8.0), flash frozen in liquid nitrogen and stored at -80 °C for further use.

Protein purification was performed by disrupting the cells by sonication with 60% amplitude and intervals of 0.5 s / 1 s pulse/pause (Bandelin Sonoplus HD 3100 homogenizer). Cellular debris was removed by centrifugation of the lysate at 30,000×f*g* for 30 min at 4 °C. All proteins were purified by gravity flow Ni^2+^ -NTA affinity chromatography (Qiagen, Cat. -No. 30230), except TEV-construct which was purified by gravity flow Strep-Tactin®XT Superflow® high-capacity resin (IBA Cat. -No.2-1208-002). Ni^2+^ -NTA purification was performed following the manufacturer’s instruction. Briefly, 1.5 mL settled beads of Ni^2+^ -NTA resin were equilibrated with 15 mL Ni Lysis buffer. Cleared lysate was loaded onto the gravity flow column, washed twice with 30 mL Ni Wash buffer (50 mM NaH_2_PO_4_, 300 mM NaCl, and 20 mM imidazole, pH 8.0, or increasing imidazole concentrations of 20 mM, 40 mM, and 60 mM imidazole, each 20 mL wash volume for TVMV) and eventually eluted with 8 mL Ni Elution buffer (50 mM NaH_2_PO_4_, 300 mM NaCl, and 250 mM imidazole, pH 8.0). For TEV-construct 500 µL bed volume of the Strep-Tactin®XT Superflow® high-capacity resin was equilibrated with 1 mL buffer W (100 mM Tris/HCl, 150 mM NaCl, pH 8.0). After applying the cleared lysate onto the resin, the column was washed twice with 2.5 mL buffer W. Proteins were finally eluted by applying buffer BXT (100 mM Tris/HCl, 150 mM NaCl, and 50 mM biotin, pH 8.0) and pooling the fractions containing the protein of interest.

All eluted proteins were supplemented with 10 mM 2-mercaptoethanol (2-ME) directly following purification and dialysed against phosphate-buffered saline (PBS: 2.7 mM KCl, 1.5 mM KH_2_PO_4_, 8.1 mM Na_2_HPO_4_, 137 mM NaCl) supplemented with 10 mM 2-ME using SnakeSkin dialysis tubing (10kDa MWCO) (Fisher Scientific, Cat. -No. 10005743). Eventually proteins were supplemented with 15% (v/v) glycerol and frozen at -80 °C until further use. Proteins were analysed by SDS-PAGE (15% (w/v) gels at 160 V and stained by Coomassie brilliant blue staining (Figure S8). Identity and integrity of the proteins was monitored by mass-spectrometry (**Figure S9** and **Table S3**). Further purification of the proteins towards achieving less impurities was not carried out, because the decoder system assembly on the beads would serve as a second and more targeted protein purification. The concentration of the proteins was determined either by Bradford assay using protein assay dye reagent (Bio-Rad) and bovine serum albumin (Carl Roth) as standard. Measurement was performed in a Multiskan GO spectrophotometer at 575 nm (Thermo Fisher Scientific). The reported concentration values correspond to the eluted protein solutions. For setting up the assays, the proteases have been quantified via their enzymatic activity.

The specific activities of the engineered split TEV and TVMV constructs were compared to the full-versions of these proteases (Figure S11).

### Material module A containing TEV

Epoxy-activated magnetic beads (Cube Biotech, Cat. -No. 50805) were functionalized with novobiocin according to manufacturer’s instructions with adjustments. Briefly, epoxy-activated material was washed 4 times using a magnetic stand with water, followed by equilibration in coupling buffer (100 mM Na_2_CO_3_, 500 mM NaCl, pH 8.3). A 125 mM solution of novobiocin in coupling buffer was mixed with the material (2 mL solution per 2 mL 25% (v/v) slurry) and incubated at 37 °C for 21 h while shaking at 150 rpm. Excess novobiocin was washed away by coupling buffer by separating the beads using a magnetic stand from the supernatant. This was followed by 5 rounds of 2 mL wash with water by adding the wash buffer, gently inverting the tube, and collecting the beads by a magnetic stand and subsequently removing the supernatant. The reaction was quenched by quenching buffer (1 M ethanolamine/HCl, pH 7.4) at 37 °C for 8 h while shaking at 150 rpm. Finally, the material was blocked with 1% (w/v) BSA in PBS overnight at 4 °C with agitation. Binding capacity of the functionalized beads was determined by adding an excess amount of GyrB-fused TEV protease, incubation overnight at 4 °C in an end-to-end rotator, and measuring the quantity of non-bound TEV protease in the supernatant by measuring the TEV activity of the supernatant and measuring its quantity using a calibration standard (6 data points of concentrations between 0 – 0.235 mg mL^-1^ TEV in PBS (supplemented with 10 mM 2-ME), 16 µL per well). We obtained a binding capacity of 7.8 mg protein per mL settled novobiocin-functionalized magnetic beads. For decoder experiments, the novobiocin-functionalized material was supplemented with a 25% excess amount of GyrB-fused TEV protease in PBS containing 10 mM 2-ME and 0.05% (v/v) Tween®20. The suspension was incubated for 16 h at 4 °C on an end-to-end rotator. Unbound protein was eliminated in 6 washing steps using 2 mL of the same buffer.

### Material module B containing TVMV

TVMV protein fused to a C-terminal His_6_-tag was mixed with spherical magnetic Ni^2+^ -NTA functionalized agarose beads (Cube Biotech, Cat. -No. 31201). The binding capacity of the material is reported as 80 mg mL^-1^ of the settled beads by the manufacturer. The binding was performed according to the product instructions. Briefly, the beads were washed twice with Ni Lysis buffer, and mixed with the His_6_-tagged TVMV protease in Ni Lysis buffer supplemented with 10 mM 2-ME and 0.05% (v/v) Tween®20 for 16 h at 4 °C on an end-to-end rotator. After coupling, unbound protein was eliminated by 6 washing steps with the same buffer.

### Material module C containing Casp3

Casp3-containing matrix consisted of an N- and C-terminally HaloTag®-ed Casp3 coupled to Magne® HaloTag beads (Promega, Cat.-No. G7281). To ensure binding on both termini, proteins with degraded C-terminal were eliminated by a pre-purification step on magnetic Ni^2+^ -NTA beads (the His_6_-tag is at the C-terminus), **Figure S10**. Briefly, the proteins were coupled onto Ni^2+^ -NTA beads in Ni Lysis buffer supplemented with 10 mM 2-ME and 0.05% (v/v) Tween®20 for 4 h at 4 °C on an end-to-end rotator. The beads were then washed 6 times with Ni Lysis buffer and finally resuspended in PBS containing 10 mM 2-ME, 0.05% (v/v) Tween®20, and 100 mM EDTA. The suspension was incubated at room temperature for 1 h while being gently mixed on an orbital shaker. Finally, the supernatant which contained the full-length protein was added to Magne® HaloTag beads prewashed with PBS containing 10 mM 2-ME and 0.005% (v/v) IGEPAL. Coupling was performed for 16 h at 4 °C on an end-to-end rotator. Unbound protein was eliminated by washing 6 times with 2 mL PBS containing mM 2-ME and 0.005 % (v/v) IGEPAL. Beads-immobilization steps of the 3 modules were carried out in 2-mL tubes and tubes were filled to avoid the drying out of the mixtures.

### System assembly

The system was assembled in 12-well plates placed on a support with 3 magnets (4 mm diameter, Supermagnete-Webcraft GmbH, cat. No S-04-10-AN) under each well. For assembling the system, each of the 3 modules (60 µg TEV-/, 30 µg TVMV-/, and 40 µg Casp3-bound material each resuspended in 50 µL PBS containing 10 mM 2-ME) were pipetted on top of one of the magnets in the 12-well-plate to a final volume of 550 µL of assay buffer (PBS supplemented with 10 mM 2-ME, 5 mM TRIS/HCl, 75 ng µL^-1^ BSA) for direct immobilization. The wells were filled with 53.6 µM novobiocin, 10 mM EDTA, or both as indicated. The system was incubated at 4 °C on a tilting shaker.

Release kinetics were carried out by taking samples from the supernatant in each well and measuring the activity of each of the output proteases in all conditions (section Analytical methods).

### Analytical methods

TEV activity was determined using the Senso-Lyte 520 TEV Activity Assay Kit according to the manufacturer’s protocol with the following changes: 8 µL of substrate solution was added to 8 µL of TEV containing buffer in a black low-volume 384-well plate directly prior to measurement. Fluorescence (excitation: 490 nm, emission: 520 nm) was measured every minute for 60 minutes. The activity was determined by calculating the slope in the linear range and by using a calibration curve with 5-FAM (0-1000 nM in PBS supplemented with 10 mM 2-ME, 16 µL per well). One relative unit (RU) for TEV was defined as the amount of cleaved substrate equivalent to the fluorescence of 1 fmol 5-FAM per min at 34 °C.

TVMV activity was determined using a custom-synthesized peptide by Genscript containing the substrate for TVMV protease (GETVRFQSDT) and a FRET pair of 7-Methoxycoumarin-4 (MCA) and 2,4-dinitrophenyl (DNP). TVMV substrate was dissolved in DMSO to a final concentration of 5 mM and stored at -80 °C. For activity measurements, 8 µL of 300 µM final concentration of substrate diluted in TVMV protease assay buffer (50 mM Tris/HCl, 100 mM NaCl, and 2 mM DTT, 50 mg mL^-1^ BSA (Sigma-Aldrich), pH 8.0) was added to 8 µL TVMV sample. The fluorescence (excitation: 325 nm, emission 392 nm) was measured every minute for 60 minutes at 34 °C. Activity was determined by measuring the slope in the linear range, using a dilution series of 0-125 nM MCA under the assay conditions. One relative unit for TVMV was defined as the amount of cleaved substrate equivalent to the fluorescence of 1 fmol MCA per min at 34 °C.

Casp3 activity was determined using the Caspase-3 Colorimetric Assay Kit according to manufacturer’s instructions with slight changes. 10 µL of the substrate diluted in PBS supplemented with 10 mM 2-ME was added to 10 µL of the samples in a transparent 384-well plate. The absorbance at 405 nm was measured every minute for 1 h at 34 °C. Activity was determined by measuring the slope in the linear range using a dilution series of p-nitroaniline (pNA) (0–1000 µM in PBS containing 10 mM 2-ME) that was used for calibration.

1 RU corresponded to the amount of cleaved substrate equivalent to 1 pmol of pNA per min at 34°C.

### Statistical Analysis

Bar charts show the mean values of replicates, and replicates are shown separately on each figure. In each data set, the negative slope of the negative control was set to zero, and all data points were normalized to this value.

## Supporting information

Supplementary Figures 1-11, Supplementary Tables 1-3, and supplementary method of the mathematical model

## Supporting Information

Supporting Information is available from the Wiley Online Library or from the author.

## Acknowledgements

This work was supported by the German Research Foundation (Deutsche Forschungsgemeinschaft, DFG) under Germany’s Excellence Strategy – CIBSS, EXC-2189, Project ID: 390939984, and through the grant TRR 179 (Grant No. 272 983 813), under the Excellence Initiative of the German Federal and State Governments – BIOSS, EXC-294, by the German Ministry of Education and Research (BMBF) through LiSyM Cancer (Grant No. 031L0256G) and in part by the Ministry for Science, Research and Arts of the State of Baden-Württemberg. The authors are thankful to Melissa Klenzendorf and Denise Gaspar for their technical support. The authors would like to sincerely thank Bettina Knapp for excellent technical assistance in the data acquisition of MS-Experiments.

Received: ((will be filled in by the editorial staff))

Revised: ((will be filled in by the editorial staff))

Published online: ((will be filled in by the editorial staff))

## References

[1] J. A. N. Brophy, C. A. Voigt, Nat. Methods 2014, 11, 508.

[2] D. E. Cameron, C. J. Bashor, J. J. Collins, Nat. Rev. Microbiol. 2014, 12, 381.

[3] Z. Chen, R. D. Kibler, A. Hunt, F. Busch, J. Pearl, M. Jia, Z. L. VanAernum, B. I. M. Wicky, G. Dods, H. Liao, M. S. Wilken, C. Ciarlo, S. Green, H. El-Samad, J. Stamatoyannopoulos, V. H. Wysocki, M. C. Jewett, S. E. Boyken, D. Baker, Science (80-.). 2020, 368, 78.

[4] X. J. Gao, L. S. Chong, M. S. Kim, M. B. Elowitz, Science (80-.). 2018, 361, 1252.

[5] A. A. K. Nielsen, B. S. Der, J. Shin, P. Vaidyanathan, V. Paralanov, E. A. Strychalski, D. Ross, D. Densmore, C. A. Voigt, Science (80-.). 2016, 352.

[6] B. Groves, Y. J. Chen, C. Zurla, S. Pochekailov, J. L. Kirschman, P. J. Santangelo, G. Seelig, Nat. Nanotechnol. 2016, 11, 287.

[7] R. Daniel, J. R. Rubens, R. Sarpeshkar, T. K. Lu, Nature 2013, 497, 619.

[8] A. A. Green, J. Kim, D. Ma, P. A. Silver, J. J. Collins, P. Yin, Nature 2017, 548, 117.

[9] Paul, E. M. Warszawik, M. Loznik, A. J. Boersma, A. Herrmann, Angew. Chemie 2020, 132, 20508.

[10] L. Schukur, B. Geering, G. Charpin-El Hamri, M. Fussenegger, Sci. Transl. Med. 2015, 7, 1.

[11] X. Wan, F. Volpetti, E. Petrova, C. French, S. J. Maerkl, B. Wang, Nat. Chem. Biol. 2019, 15, 540.

[12] A. Tamsir, J. J. Tabor, C. A. Voigt, Nature 2011, 469, 212.

[13] T. Fink, J. Lonzaric, A. Praznik, T. Plaper, E. Merljak, K. Leben, N. Jerala, T. Lebar, Ž. Strmšek, F. Lapenta, M. Benčina, R. Jerala, Nat. Chem. Biol. 2019, 15, 115.

[14] H. J. Wagner, R. Engesser, K. Ermes, C. Geraths, J. Timmer, W. Weber, Mater. Today 2019, 22, 25.

[15] H. J. Wagner, S. Kemmer, R. Engesser, J. Timmer, W. Weber, Adv. Sci. 2019, 6.

[16] H. M. Beyer, R. Engesser, M. Hörner, J. Koschmieder, P. Beyer, J. Timmer, M. D. Zurbriggen, W. Weber, Adv. Mater. 2018, 30, 1.

[17] P. M. Gawade, J. A. Shadish, B. A. Badeau, C. A. DeForest, Adv. Mater. 2019, 31, 1.

[18] P. Zhang, D. Gao, K. An, Q. Shen, C. Wang, Y. Zhang, X. Pan, X. Chen, Y. Lyv, C. Cui, T. Liang, X. Duan, J. Liu, T. Yang, X. Hu, J. J. Zhu, F. Xu, W. Tan, Nat. Chem. 2020, 12, 381.

[19] H. Komatsu, S. Matsumoto, S. ichi Tamaru, K. Kaneko, M. Ikeda, I. Hamachi, J. Am. Chem. Soc. 2009, 131, 5580.

[20] M. Ikeda, T. Tanida, T. Yoshii, K. Kurotani, S. Onogi, K. Urayama, I. Hamachi, Nat. Chem. 2014, 6, 511.

[21] L. Liu, J. A. Shadish, C. K. Arakawa, K. Shi, J. Davis, C. A. DeForest, Adv. Biosyst. 2018, 2, 1.

[22] S. T. Koshy, T. C. Ferrante, S. A. Lewin, D. J. Mooney, Biomaterials 2014, 35, 2477.

[23] A. P. Liu, E. A. Appel, P. D. Ashby, B. M. Baker, E. Franco, L. Gu, K. Haynes, N. S. Joshi, A. M. Kloxin, P. H. J. Kouwer, J. Mittal, L. Morsut, V. Noireaux, S. Parekh, R. Schulman, S. K. Y. Tang, M. T. Valentine, S. L. Vega, W. Weber, N. Stephanopoulos, O. Chaudhuri, Nat. Mater. 2022, 21, 390.

[24] D. Ausländer, S. Ausländer, X. Pierrat, L. Hellmann, L. Rachid, M. Fussenegger, Nat. Methods 2018, 15, 57.

[25] A. Badeau, M. P. Comerford, C. K. Arakawa, J. A. Shadish, C. A. Deforest, Nat. Chem. 2018, 10, 251.

[26] H. Faris, M. Habib, M. Faris, M. Alomari, A. Alomari, J. Biomed. Inform. 2020, 109, 103525.

[27] K. Chen, R. Wang, J. Huang, F. Gao, Z. Yuan, Y. Qi, H. Wu, Sci. Data 2022, 9, 1.

[28] N. Naseer, M. J. Hong, K. S. Hong, Exp. Brain Res. 2014, 232, 555.

[29] V. Stein, M. Nabi, K. Alexandrov, ACS Synth. Biol. 2017, 6, 1337.

[30] A. Praznik, T. Fink, N. Franko, J. Lonzaric, M. Benčina, N. Jerala, T. Plaper, S. Roškar, R. Jerala, Nat. Commun. 2022, 13, 1.

[31] H. K. Chung, M. Z. Lin, Nat. Methods 2020, 17, 885.

[32] A. Gräwe, J. Ranglack, W. Weber, V. Stein, Curr. Opin. Biotechnol. 2020, 63, 1.

[33] D. Lee, M. Creed, K. Jung, T. Stefanelli, D. J. Wendler, W. C. Oh, N. L. Mignocchi, C. Lüscher, H. B. Kwon, Nat. Methods 2017, 14, 495.

[34] V. Siciliano, B. Diandreth, B. Monel, J. Beal, J. Huh, K. L. Clayton, L. Wroblewska, A. McKeon, B. D. Walker, R. Weiss, Nat. Commun. 2018, 9, 1.

[35] T. Fink, R. Jerala, Curr. Opin. Chem. Biol. 2022, 68, 102146.

[36] V. Stein, K. Alexandrov, Trends Biotechnol. 2015, 33, 101.

[37] M. C. Wehr, L. Reinecke, A. Botvinnik, M. J. Rossner, BMC Biotechnol. 2008, 8, 1.

[38] H. Gradišar, R. Jerala, J. Pept. Sci. 2011, 17, 100.

[39] Lainšček, V. Forstnerič, V. Mikolič, Š. Malenšek, P. Pečan, M. Benčina, M. Sever, H. Podgornik, R. Jerala, Nat. Commun. 2022, 13.

[40] Lapenta, J. Aupič, M. Vezzoli, Ž. Strmšek, S. Da Vela, D. I. Svergun, J. M. Carazo, R. Melero, R. Jerala, Nat. Commun. 2021, 12, 1.

[41] T. Lebar, D. Lainšček, E. Merljak, J. Aupič, R. Jerala, Nat. Chem. Biol. 2020, 16, 513.

[42] Budihardjo, H. Oliver, M. Lutter, X. Luo, X. Wang, Annu. Rev. Cell Dev. Biol. 1999, 15, 269.

[43] D. C. Gray, S. Mahrus, J. A. Wells, Cell 2010, 142, 637.

[44] H. J. Wagner, W. Weber, Molecules 2019, 24.

[45] N. Gurdo, D. C. Volke, P. I. Nikel, Trends Biotechnol. 2022, 40, 1148.

[46] P. Carbonell, A. J. Jervis, C. J. Robinson, C. Yan, M. Dunstan, N. Swainston, M. Vinaixa, K. A. Hollywood, A. Currin, N. J. W. Rattray, S. Taylor, R. Spiess, R. Sung, A. R. Williams, D. Fellows, N. J. Stanford, P. Mulherin, R. Le Feuvre, P. Barran, R. Goodacre, N. J. Turner, C. Goble, G. G. Chen, D. B. Kell, J. Micklefield, R. Breitling, Takano, J. L. Faulon, N. S. Scrutton, Commun. Biol. 2018, 1, 1.

[47] N. A. Gormley, G. Orphanides, A. Meyer, P. M. Cullis, A. Maxwell, Biochemistry 1996, 35, 5083.

[48] S. Knecht, D. Ricklin, A. N. Eberle, B. Ernst, J. Mol. Recognit. 2009, 22, 270.

[49] C. Geraths, E. H. Christen, W. Weber, Macromol. Rapid Commun. 2012, 33, 2103.

[50] C. Geraths, M. Daoud-El Baba, G. Charpin-El Hamri, W. Weber, J. Control. Release 2013, 171, 57.

[51] M. Ehrbar, R. Schoenmakers, E. H. Christen, M. Fussenegger, W. Weber, Nat. Mater. 2008, 7, 800.

[52] O. S. Thomas, B. Rebmann, M. Tonn, I. C. Schirmeister, S. Wehrle, J. Becker, G. J. Zea Jimenez, S. Hook, S. Jäger, M. Klenzendorf, M. Laskowski, A. Kaier, G. Pütz, M. D. Zurbriggen, W. Weber, M. Hörner, H. J. Wagner, Small 2022, 18.

[53] E. H. Christen, M. Karlsson, M. M. Kämpf, R. Schoenmakers, R. J. Gübeli, H. M. Wischhusen, C. Friedrich, M. Fussenegger, W. Weber, Adv. Funct. Mater. 2011, 21, 2861.

[54] M. Hörner, K. Raute, B. Hummel, J. Madl, G. Creusen, O. S. Thomas, E. H. Christen, N. Hotz, R. J. Gübeli, R. Engesser, B. Rebmann, J. Lauer, B. Rolauffs, J. Timmer, W. W. A. Schamel, J. Pruszak, W. Römer, M. D. Zurbriggen, C. Friedrich, A. Walther, S. Minguet, R. Sawarkar, W. Weber, Adv. Mater. 2019, 31, 1.

[55] X. Zhang, C. Dong, W. Huang, H. Wang, L. Wang, D. Ding, H. Zhou, J. Long, T. Wang, Z. Yang, Nanoscale 2015, 7, 16666.

[56] E. Chervyachkova, S. V. Wegner, ACS Synth. Biol. 2018, 7, 1817.

[57] M. A. English, L. R. Soenksen, R. V Gayet, H. De Puig, N. M. Angenent-mari, A. S. Mao, P. Q. Nguyen, J. J. Collins, 2019, 785, 780.

[58] M. M. Kaminski, O. O. Abudayyeh, J. S. Gootenberg, F. Zhang, J. J. Collins, Nat. Biomed. Eng. 2021, 5, 643.

[59] F. Cella, L. Wroblewska, R. Weiss, V. Siciliano, Nat. Commun. 2018, 9.

[60] E. M. Bressler, W. W. Wong, S. Adams, Y. L. Colson, M. W. Grinstaff, W. W. Wong, Y. L. Colson, 2023.

[61] J. R. Sempionatto, J. A. Lasalde-Ramírez, K. Mahato, J. Wang, W. Gao, Nat. Rev. Chem. 2022, 6, 899.

[62] H. Teymourian, C. Moonla, F. Tehrani, E. Vargas, R. Aghavali, A. Barfidokht, T. Tangkuaram, P. P. Mercier, E. Dassau, J. Wang, Anal. Chem. 2020, 92, 2291.

[63] D. G. Gibson, L. Young, R. Y. Chuang, J. C. Venter, C. A. Hutchison, H. O. Smith, Nat. Methods 2009, 6, 343.

