## Supplementary Figures 1-11, Supplementary Tables 1-3, and supplementary method of the mathematical model for "Design of a Biohybrid Materials Circuit with Binary Decoder Functionality"

Contents:

Supplementary Figures 1-11
Supplementary Tables 1-3
Supplementary Method: Description of the mathematical model (Model summary)

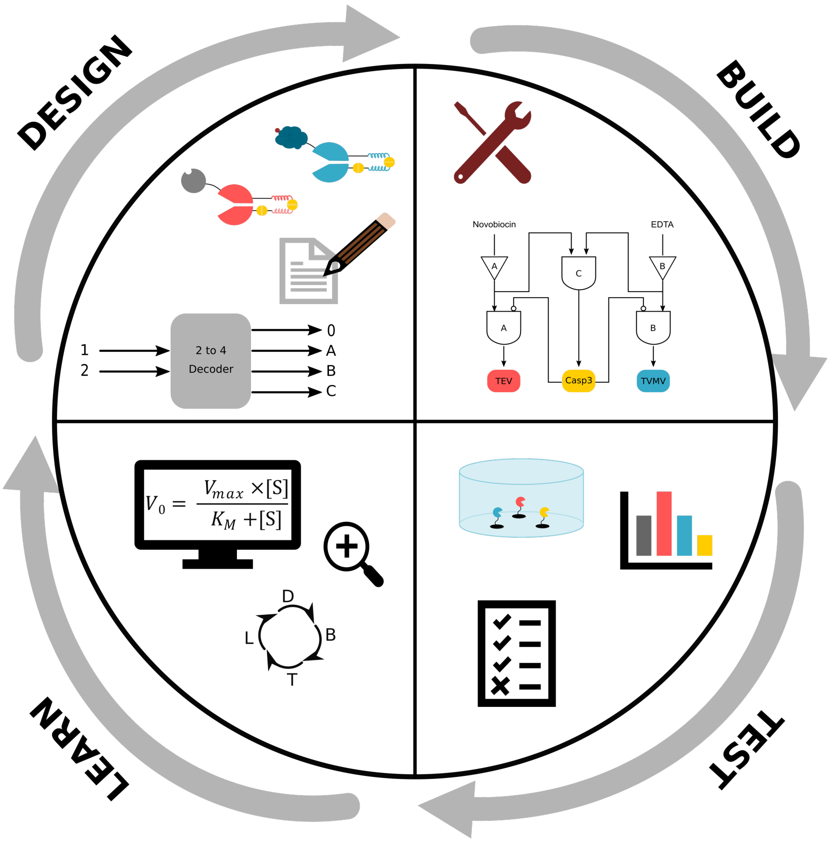


**Figure S1**. Concept of model-guided DBTL in the generation and optimization of a binary decoder. **Design**: The individual modules as well as their interplay are designed. **Build**: The decoder system is physically assembled as designed in the previous step. **Test**: The functionality of the system is tested, and data is collected. **Learn**: Collected data is extensively analysed by mathematical model-supported characterization that guides the next iteration of DBTL.

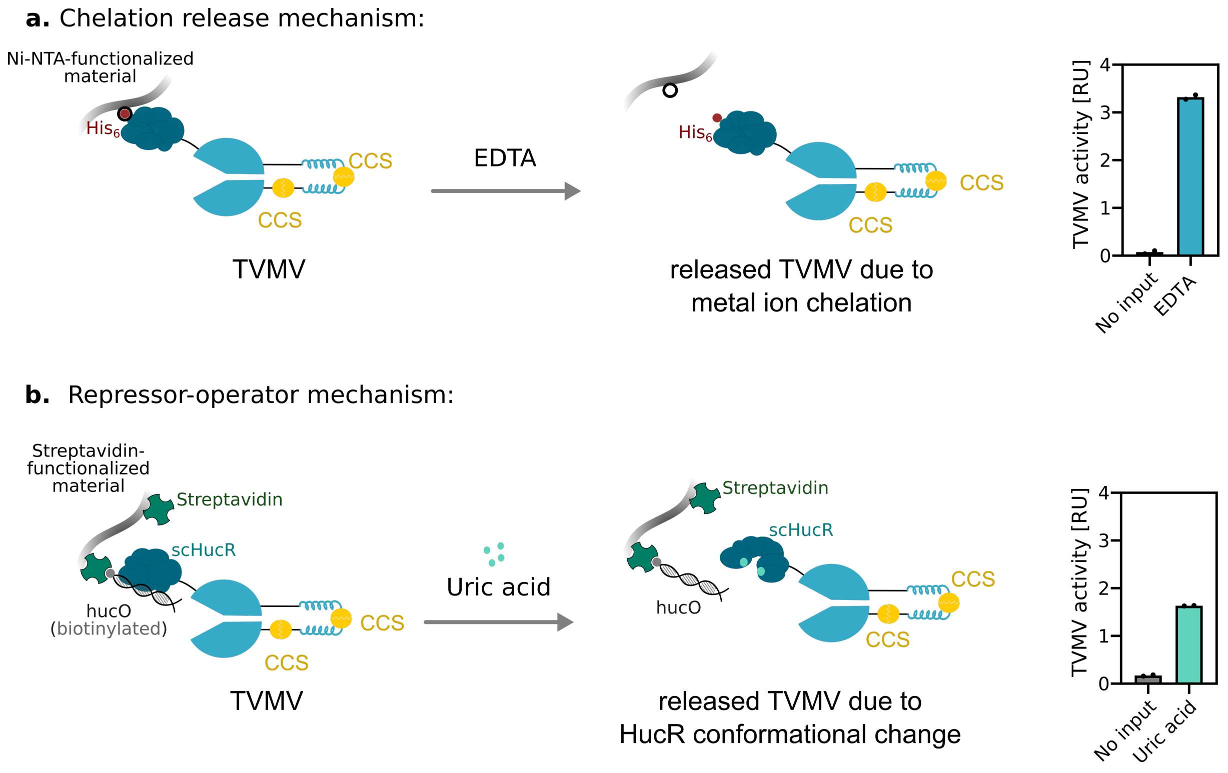


**Figure S2**. Release of TVMV protease from material modules via two mechanisms. a) Release via chelation of metal ions. 25 µg $His_6$-tagged TVMV fused to a single-chain variant of the uric acid-responsive repressor protein HucR (scHucR) was immobilized on $Ni^{2+}$-NTA-functionalized material support overnight at 4 °C prior to the addition of 10 mM EDTA. Released TVMV activity was quantified after 120 min at RT. b) TVMV-scHucR was bound to its cognate hucO operator DNA.[1] The hucO DNA sequence was immobilized via biotin on streptavidin-functionalized material support. Upon addition of uric acid, HucR undergoes a change in conformation and detaches from hucO. 25 µg of TVMV-scHucR was incubated with hucO (scHucR : hucO molar ratio of 1 : 4) for 30 min at RT prior to immobilization on streptavidin-functionalized magnetic material support for 16 h at 4 °C. Release of TVMV activity in the presence or absence of 1 mM uric acid was quantified after 120 min at RT.

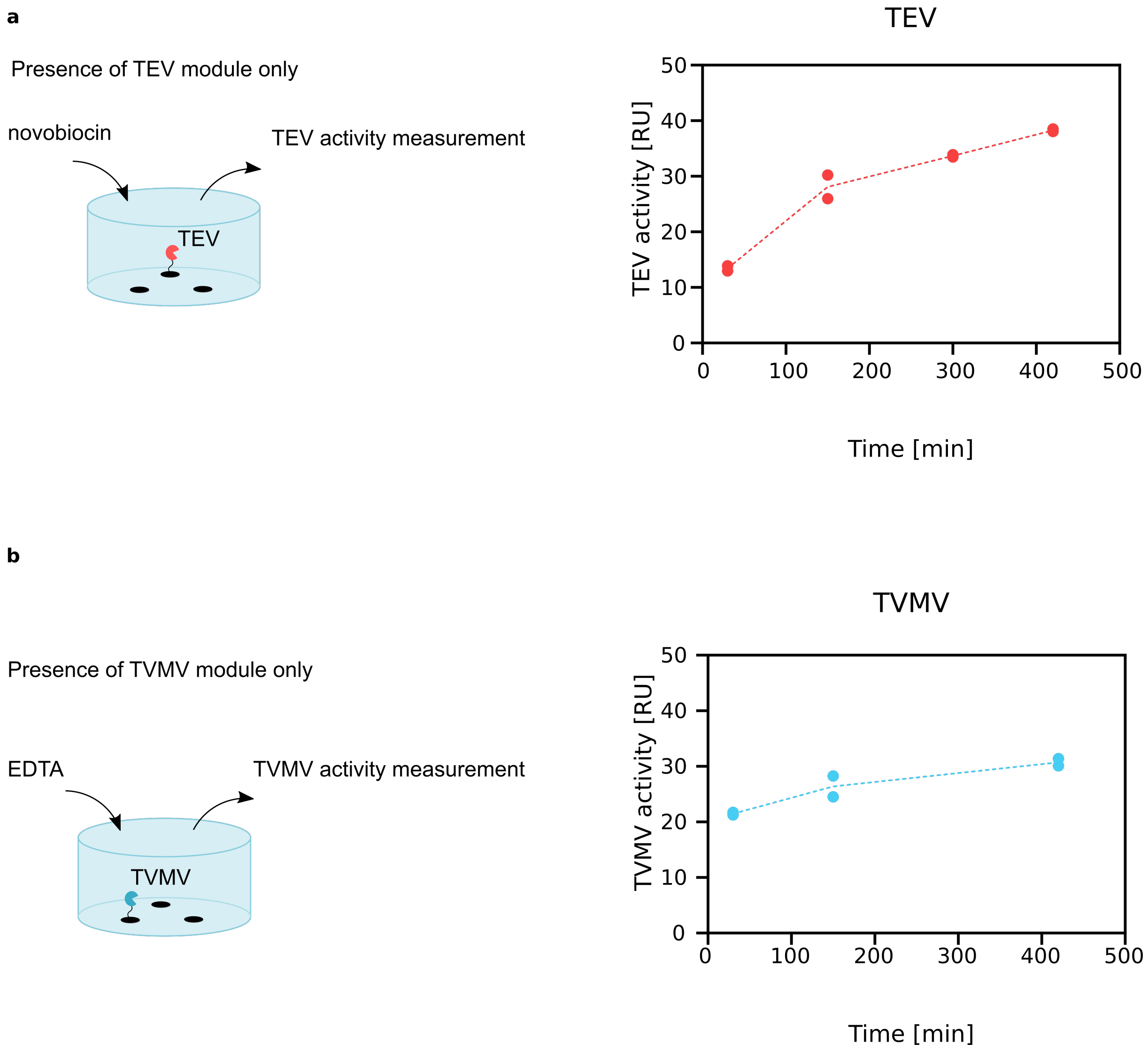


**Figure S3**. Release of TEV and TVMV protease from material modules A and B. a) Novobiocin-responsive release of TEV. Module A (60 µg TEV immobilized on its material support) was incubated in the presence of 53.6 µM novobiocin and the released TEV was quantified at the indicated points in time. b) EDTA-responsive release of TVMV. Module B (30 µg TVMV immobilized on its material support) was incubated in the presence of 10 mM EDTA and the released TVMV was quantified at the indicated points in time. RU, relative unit. Dotted lines connect the average of the replicates at each time point.

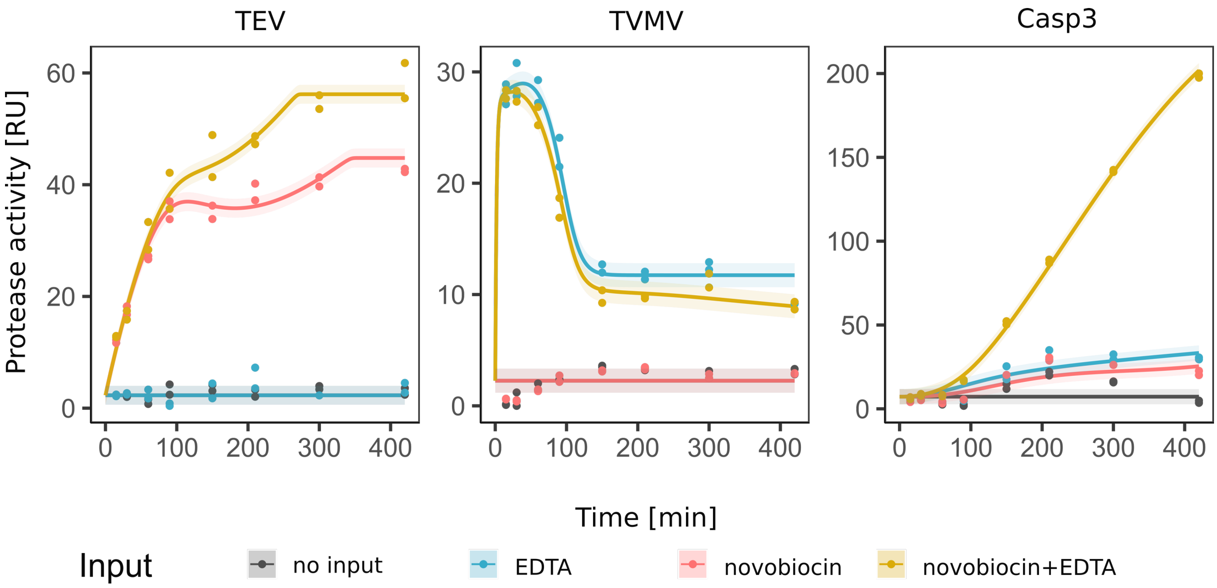


**Figure S4**. Kinetic characterization of the initial setup for the decoder circuit. The reaction was performed as described in Figure 3b. The activities of the three proteases were determined at the indicated points in time. The curves represent the model fit and the corresponding error bands. RU, relative unit.

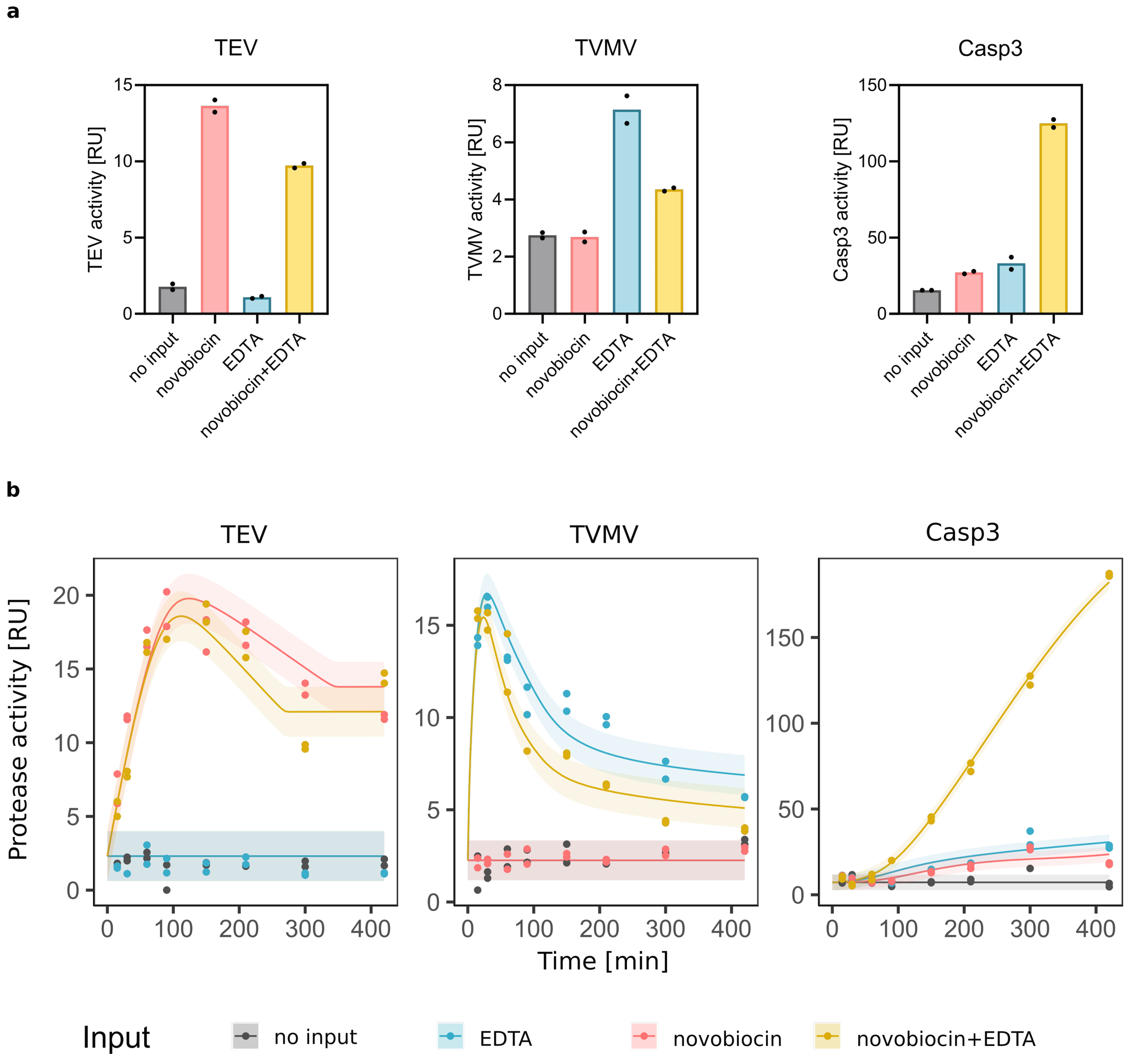


**Figure S5.** Performance of the decoder circuit with initial integration of displacer molecule treatment. a) The decoder was set up as described in Figure 3 and incubated for 300 min followed by a 30 min incubation in the presence of 0.054 mg $ml^{-1}$ of the displacer module. Bar charts indicate the mean of two separate biological replicates for each condition. b) Time resolved data corresponding to Figure S3a. At the indicated points in time, samples were taken and incubated in the presence of 0.054 mg $ml^{-1}$ displacer for 30 min prior to scoring protease activity. The curves represent the model fit to the data and the corresponding error bands.

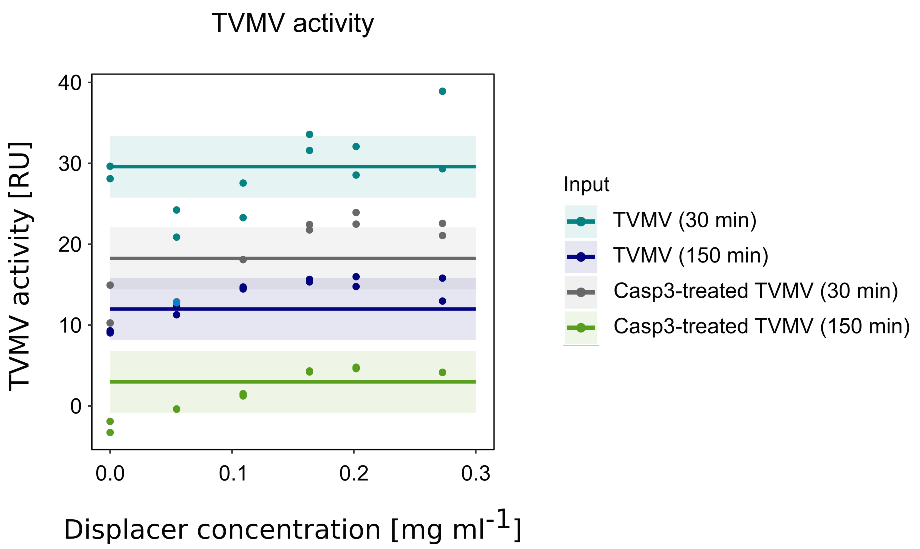


**Figure S6.** Influence of the displacer module on TVMV activity. 15 µg of TVMV was incubated in the presence of absence of 3 µg Casp3 in 300 µL assay buffer in the presence of the indicated displacer concentrations for the indicated periods of time prior to quantifying TVMV activity. In order to be coherent with the complete decoder setup, the buffer further contained 10 mM EDTA, and the buffer for Casp3-treated conditions was further supplemented with 30 µg TEV, 53.6 µM novobiocin, and 10 mM EDTA.

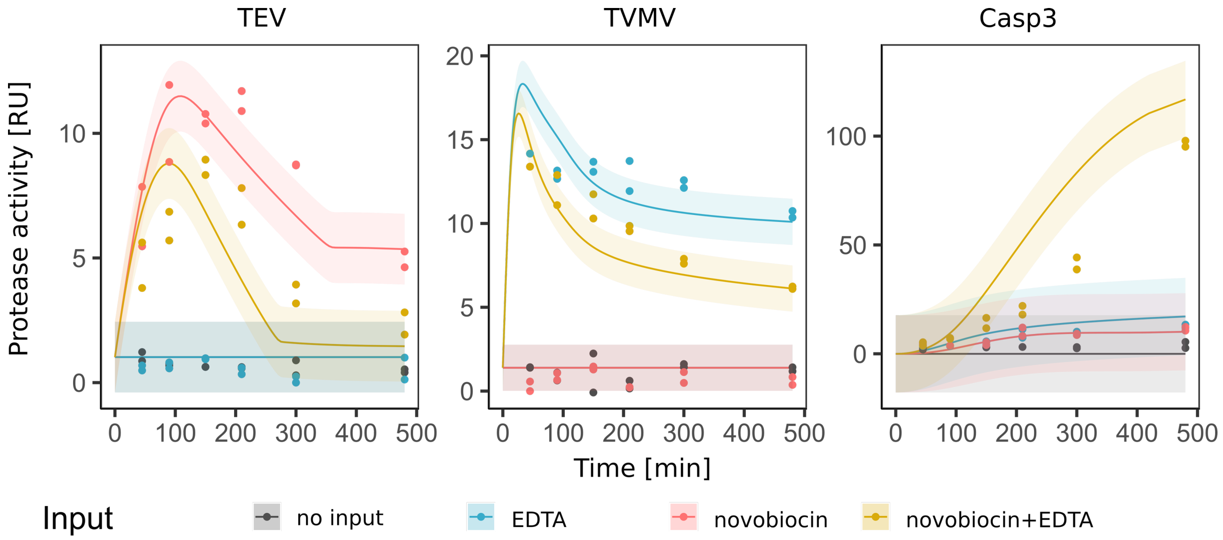


**Figure S7.** Performance of the decoder circuit with optimized displacer conditions. The experiment was performed as described in Figure 4e. At the indicated points in time, samples were taken and incubated in the presence of 0.087 mg ml$^{-1}$ displacer for 2.5 h prior to quantifying the activities of the three proteases. The curves and the error band represent the model predictions.

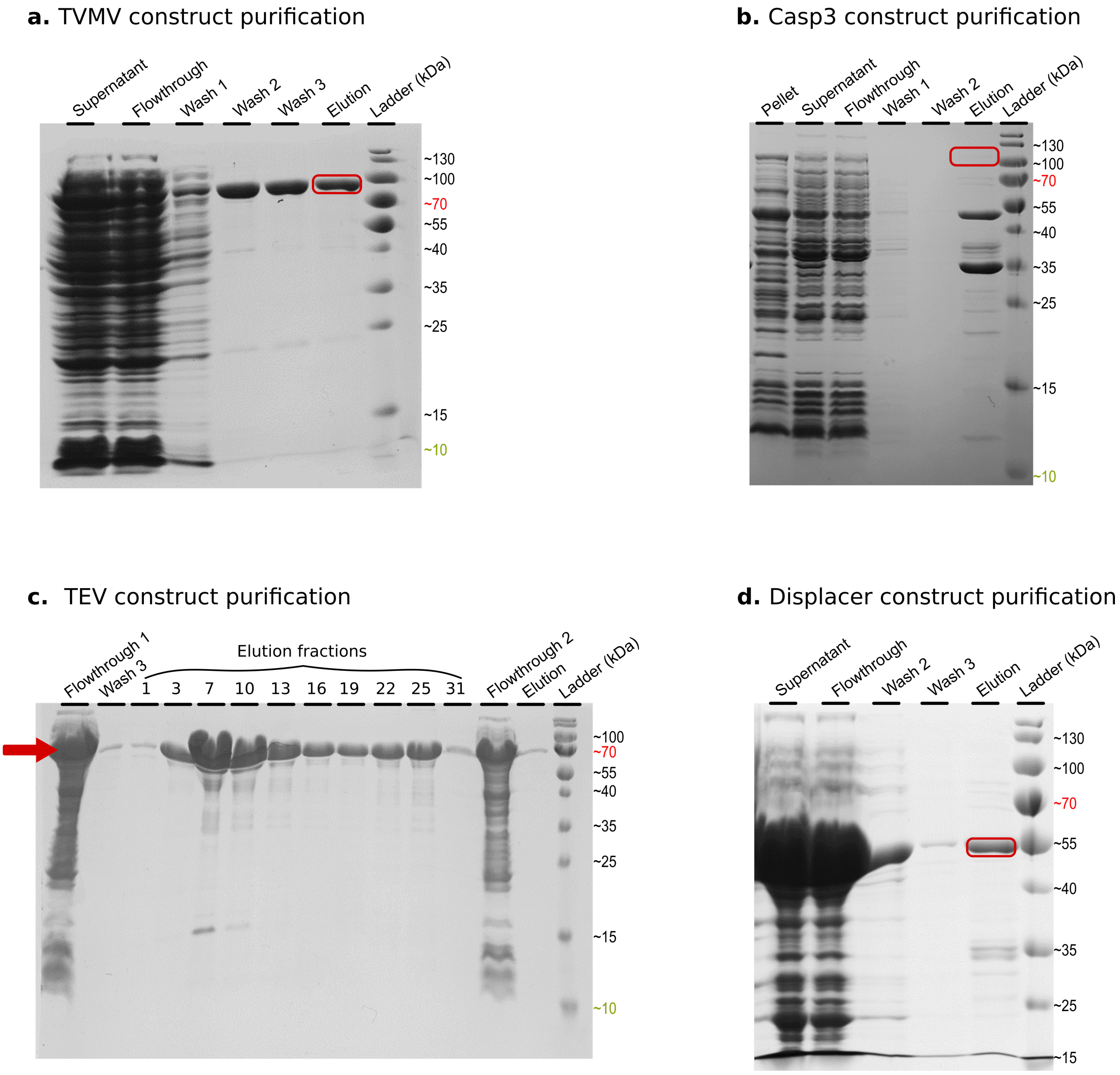


**Figure S8**. SDS-PAGE analysis of the purification of protein constructs. a) Analysis of TVMV construct purification via IMAC. Wash steps contain increasing imidazole concentrations of 20 mM, 40 mM, and 60 mM. Expected size of the protein is 77.2 kDa. b) Analysis of Casp3 construct purification via IMAC and two wash steps, both containing 20 mM imidazole as described in the methods section. Expected size of the construct is 105.8 kDa. c) Analysis of TEV construct purification by gravity flow Strep-Tactin®XT Superflow® high-capacity resinas described in the methods section. Elution fractions of 500 µL were collected step-wise and subsequently mixed together. Flowthrough of the first purification run was purified for another round by fresh Strep-Tactin resin. Expected protein size is 64 kDa. d) Purification of displacer construct via IMAC. Protein is expected to have a size of 47.6 kDa. All gels were run on a 15% (w/v) SDS-gel.

**a.** pHM118 construct

1 MGSAWSHPQF EKGGGSGGGS GGSAWSHPQF EKSAGKSMSS MVSDTSCTFP
51 SSDGIFWKHW IQTKDGQCGS PLVSTRDGFI VGIHSASNFT NTNNYFTSVP
101 KNFMELLTNQ EAQQWVSGWR LNADSVLWGG HKVFMVGGGS GGGDEVDGGG
151 SGGGSPEDEL AANEEELQQN EQKLAQIKQK LQAIKYGGGS GGDEVDGGSG
201 GSPEDEIQQL EEEIAQLEQK NAALKEKNQA LKYGGSGGSG GSGGSGGSGG
251 GESLFKGPRD YNPISSTICH LTNESDGHTT SLYGIGFGPF IITNKHLFRR
301 NNGTLLVQSL HGVFKVKNTT TLQQHLIDGR DMIIIRMPKD FPPFPQKLKF
351 REPQREERIC LVTTNFQTGS GSGSGSGSGS NSYDSSSIKV LKGLDAVRKR
401 PGMYIGDTDD GTGLHHMVFE VVDNAIDEAL AGHCKEIIVT IHADNSVSVQ
451 DDGRGIPTGI HPEEGVSAAE VIMTVLHAGG KFDDNSYKVS GGLHGVGVSV
501 VNALSQKLEL VIQREGKIHR QIYEHGVPQA PLAVTGETEK TGTMVRFWPS
551 LETFTNVTEF EYEILAKRLR ELSFLNSGVS IRLRDKRDGK EDHFHYEG

Split-TEV
(99% sequence coverage)

GyrB
(95% sequence coverage)

**b.** pHM83 construct

1 MKSVSSLVSE SSHIVHKEDT SFWQHWITTK DGQCGSPLVS IIDGNILGIH
51 SLTHTTNGSN YFVEFPEKFV ATYLDAADGW CKNWKFNADK ISWGSFTLVE
101 DAPEDGGGSG GGDEVDGGGS GGGSPEDENA QLEQKNAQLK QEISQLEQEI
151 GGSGGDEVDG GSGGSPEDKL AQIKEKLQQI KEELAANEEK LQANKYGGGG
201 SGGGSGGGSG GGSSKALLKG VRDFNPISAC VCLLENSSDG HSERLFGIGF
251 GPYIIANQHL FRRNNGELTI KTMHGEFKVK NSTQLQMKPV EGRDIIVIKM
301 AKDFPPFPQK LKFRQPTIKD RVCMVSTNFQ QGGGGSGGGG SGGGGSARMD
351 NDTAALLERI RSDWARLNHG QGPDSDGLTP SAGPMLTLLL LERLHAALGR
401 EIERTYAASG LNAAGWDLLL TLYRSAPPEG LRPTELSALA AISGPSTSNR
451 IVRLLEKGLI ERREDERDRR SASIRLTPQG RALVTHLLPA HLATTQRVLA
501 PLSAQEQRTL EELAGRMLAG LEQGVGSGGA RMSARMDNDT AALLERIRSD
551 WARLNHGQGP DSDGLTPSAG PMLTLLLLER LHAALGREIE RTYAASGLNA
601 AGWDLLLTLY RSAPPEGLRP TELSALAAIS GPSTSNRIVR LLEKGLIERR
651 EDERDRRSAS IRLTPQGRAL VTHLLPAHLA TTQRVLAPLS AQEQRTLEEL
701 AGRMLAGLEQ GVGGSGGHHH HHH

Split-TVMV
(100% sequence coverage)

scHucR
(19% sequence coverage)

**b.** pHM111 construct

1 MEIGTGFPFD PHYVEVLGER MHYVDVGPRD GTPVLFLHGN PTSSYVWRNI
51 IPHVAPTHRC IAPDLIGMGK SDKPDLGYFF DDHVRFMDAF IEALGLEEVV
101 LVIHDWGSAL GFHWAKRNPE RVKGIAFMEF IRPIPTWDEW PEFARETFQA
151 FRTTDVGRKL IIDQNVFIEG TLPMGVVRPL TEVEMDHYRE PFLNPVDREP
201 LWRFPNELPI AGEPANIVAL VEEYMDWLHQ SPVPKLLFWG TPGVLIPPAE
251 AARLAKSLPN CKAVDIGPGL NLLQEDNPDL IGSEIARWLS TLEISGHGGG
301 SGGGSGGETV RFQSGGGPAG EASSIPNREG KPIPNPLLGL GSTRTGESGI
351 SLDNSYKMDY PEMGLCIIIN NKNFHKSTGM TSRSGTDVDA ANLRETFRNL
401 KYEVRNKNDL TREEIVELMR DVSKEDHSKR SSFVCVLLSH GEEGIIFGTN
451 GPVDLKKITN FFRGDRCRSL TGKPKLFIIQ ACRGTELDCG IETDSGVDDD
501 MACHKIPVEA DFLYAYSTAP GYYSWRNSKD GSWFIQSLCA MLKQYADKLE
551 FMHILTRVNR KVATEFESFS FDATFHAKKQ IPCIVSMLTK ELYFYSGGGS
601 GGGGENLYFQ SGGGPAGEAS SIPNREGKPI PNPLLGLGST RTGEFEIGTG
651 FPFDPHYVEV LGERMHYVDV GPRDGTPVLF LHGNPTSSYV WRNIIPHVAP
701 THRCIAPDLI GMGKSDKPDL GYFFDDHVRF MDAFIEALGL EEVVLVIHDW
751 GSALGFHWAK RNPERVKGIA FMEFIRPIPT WDEWPEFARE TFQAFRTTDV
801 GRKLIIDQNV FIEGTLPMGV VRPLTEVEMD HYREPFLNPV DREPLWRFPN
851 ELPIAGEPAN IVALVEEYMD WLHQSPVPKL LFWGTPGVLI PPAEAARLAK
901 SLPNCKAVDI GPGLNLLQED NPDLIGSEIA RWLSTLEISG HHHHHH

Halo-Tag
(66% sequence coverage)

Casp3
(90% sequence coverage)

Halo-Tag
(10% sequence coverage)

**Figure S9.** Liquid chromatography-tandem mass spectrometry (LC-MS/MS) analysis of the three protease constructs involved in the biomolecular decoder. Samples were separated by SDS-PAGE. Visualization of proteins was carried out with colloidal Coomassie Blue, gel bands were excised, and proteins were in-gel digested using trypsin for subsequent MS analysis essentially as described previously.[3] Briefly, peptides mixtures were separated on an Ultimate 3,000 RSLCnano coupled to an Orbitrap Elite mass spectrometer (Thermo Fisher Scientific, Dreieich, Germany). Peptides were washed and concentrated on nanoEase M/Z C18 pre-columns and analyzed on a HSS C18 analytical column (both Waters Corporation, Milford, USA) using a 60 min gradient of solvent A (0.1% FA) and solvent B (50% MeOH; 30% ACN; 0.1% FA). Data dependent acquisition consisted of full MS scans in the range of m/z 370–1,700; resolution of 120,000 at m/z 400; and fragmentation of the 15 most abundant multiply charged precursor ions by collision induced dissociation. MS raw data were searched using MaxQuant (version 2.0.2.0; Tyanova et al., 2016) against the protein sequences of the protease constructs plus the *E. coli*-specific database from UniProt (release 2012_03). Proteins were identified with at least one unique peptide and a false discovery rate of 0.01 on both peptide and protein level. For estimates of relative quantification, iBAQ (intensity-based absolute quantification) intensities were used. Sequence coverages were confirmed by performing similar searches with the Mascot search engine (version 2.6, Matrix Science, London).

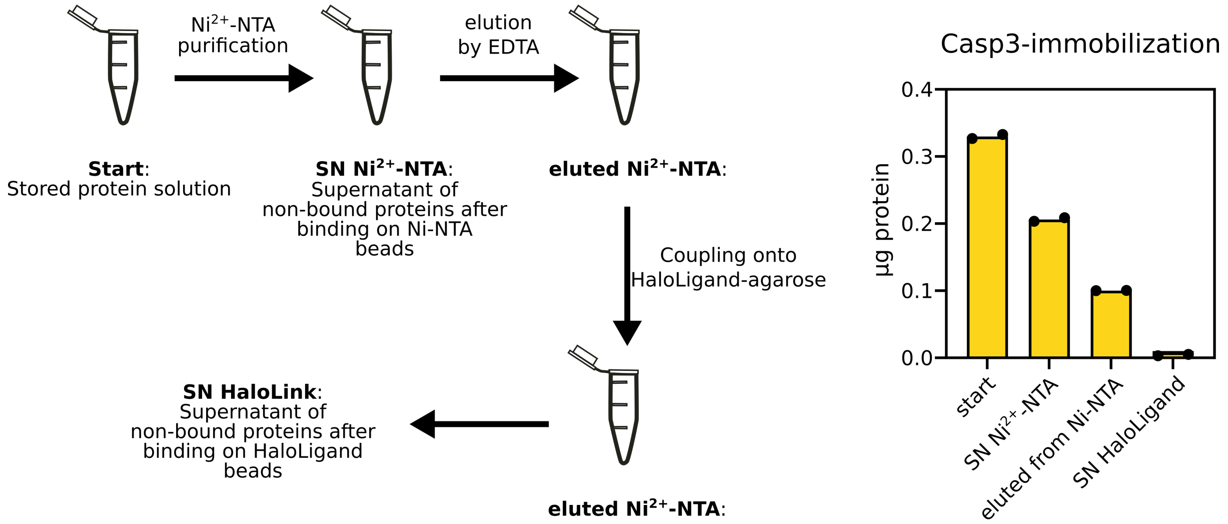


**Figure S10.** Step-wise Casp3 immobilization on its material module. Due to the low purity of Casp3 protease and the degradation of its C-terminal upon storage, immobilization of Casp3-module contained the shown steps as stated in the methods section: 320 µg of the stored protein was coupled onto $Ni^{2+}$-NTA magnetic beads, incubated for 4 h at 4 °C. Subsequently the supernatant containing the non-bound proteins was collected and stored at 4 °C. The bound proteins were subsequently eluted from beads by 100 mM EDTA, eluted samples were collected and stored at 4 °C as well. The eluted protein was then bound to HaloTag magnetic beads and incubated for 16 h at 4 °C. Samples from the supernatant after this step were acquired as well. The activity of Casp3 was measured in all samples as stated in methods section.

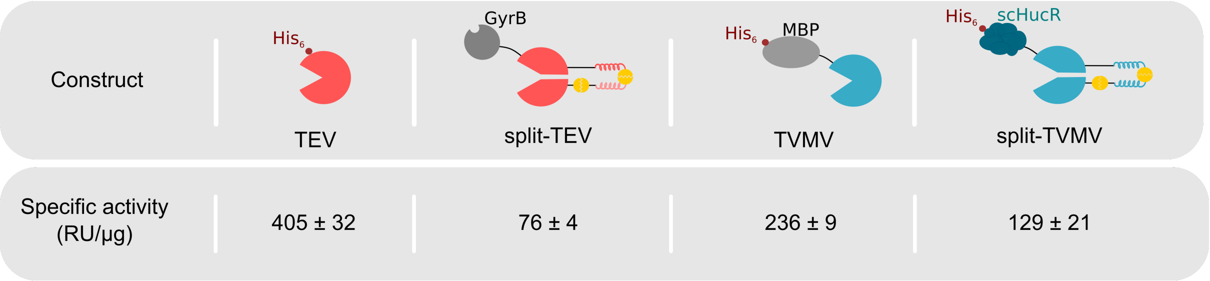


**Figure S11.** Specific activities of native and split versions of TEV and TVMV used in this work. The activity of 1.88 µg engineered split-TEV and 2.16 µg of full version TEV protease (fused to MBP as solubility tag) was determined as in the decoder experiments at 34 °C and relative enzyme units were measured as described in the methods section. Specific activities were calculated by dividing the measured activity by the mass (in µg) of protease in the system. The activity of 2 µg of each of TVMV protease versions was measured to report the relative enzyme units, and subsequently divided by the protease amounts to achieve their specific activities (RU/µg).

**Table S1.** Amino acid sequences and description of the constructs.

| Plasmid | Amino acid sequence | Description | Reference |
|---|---|---|---|
| **pHM41**<br><br>**(Displacer)** | MSPEDKIAQLKQKIQALKQENQQLEEENAALEYGGSGGSGGSGGSGGSGGKSMSSMVSDTSCTFPSSDGIFWKHWIQTKDGQAGSPLVSTRDGFIVGIHSASNFTNTNNYFTSVPKNFMELLTNQEAQQWVSGWRLNADSVLWGGHKVFMVKPEEPFQPVKEATQLMSELVYSQYPYDVPDYAGGGGSGGGGSGGGGSLEVLFQGPGSGSGSGSGSSNSYDSSSIKVLKGLDAVRKRPGMYIGDTDDGTGLHHMVFEVVDNAIDEALAGHCKEIIVTIHADNSVSVQDDGRGIPTGIHPEEGVSAAEVIMTVLHAGGKFDDNSYKVSGGLHGVGVSVVNALSQKLELVIQREGKIHRQIYEHGVPQAPLAVTGETEKTGTMVRFWPSLETFTNVTEFEYEILAKRLRELSFLNSGVSIRLRDKRDGKEDHFHYEGHHHHHH<br>P4/cTEV*/GyrB/His$_6$-Tag | Amino acid sequence for P4 coil region fused to the inactive cTEV and GyrB and His$_6$-Tag. | This work |
| **pHM83**<br><br>**(split-TVMV)** | MKSVSSLVSESSHIVHKEDTSFWQHWITTKDGQCGSPLVSIIDGNILGIHSLTHTTNGSNYFVEFPEKFVATYLDAADGWCKNWKFNADKISWGSFTLVEDAPEDGGGSGGGDEVDGGGSGGGSPEDENAQLEQKNAQLKQEISQLEQEIGGSGGDEVDGGSGGSPEDKLAQIKEKLQQIKEELAANEEKLQANKYGGGGSGGGSGGGSGGGSSKALLKGVRDFNPISACVCLLENSSDGHSERLFGIGFGPYIIANQHLFRRNNGELTIKTMHGEFKVKNSTQLQMKPVEGRDIIVIKMAKDFPPFPQKLKFRQPTIKDRVCMVSTNFQQGGGGSGGGGSGGGGSARMDNDTAALLERIRSDWARLNHGQGPDSDGLTPSAGPMLTLLLLERLHAALGREIERTYAASGLNAAGWDLLLTLYRSAPPEGLRPTELSALAAISGPSTSNRIVRLLEKGLIERREDERDRRSASIRLTPQGRALVTHLLPAHLATTQRVLAPLSAQEQRTLEELAGRMLAGLEQGVGSGGARMSARMDNDTAALLERIRSDWARLNHGQGPDSDGLTPSAGPMLTLLLLERLHAALGREIERTYAASGLNAAGWDLLLTLYRSAPPEGLRPTELSALAAISGPSTSNRIVRLLEKGLIERREDERDRRSASIRLTPQGRALVTHLLPAHLATTQRVLAPLSAQEQRTLEELAGRMLAGLEQGVGGSGGHHHHHH<br>cTVMV/CCS/P9mS/CCS/AP10/nTVMV/scHucR/His$_6$-Tag | Amino acid sequence for C-terminal split-TVMV fused to P9mS coiled region via a linker containing CCS, followed by another CCS-containing linker fusing AP10 coiled region to N-terminal of split-TVMV with a C-terminal His$_6$-Tag. | This work |
| **pHM111**<br><br>**(Casp3)** | MEIGTGFPFDPHYVEVLGERMHYVDVGPRDGTPVLFLHGNPTSSYVWRNIIPHVAPTHRCIAPDLIGMGKSDKPDLGYFFDDHVRFMDAFIEALGLEEVVLVIHDWGSALGFHWAKRNPERVKGIAFMEFIRPIPTWDEWPEFARETFQAFRTTDVGRKLIIDQNVFIEGTLPMGVVRPLTEVEMDHYREPFLNPVDREPLWRFPNELPIAGEPANIVALVEEYMDWLHQSPVPKLLFWGTPGVLIPPAEAARLAKSLPNCKAVDIGPGLNLLQEDNPDLIGSEIARWLSTLEISGHGGGSGGGSGGETVRFQSGGGPAGEASSIPNREGKPIPNPLLGLGSTRTGESGISLDNSYKMDYPEMGLCIIINNKNFHKSTGMTSRSGTDVDAANLRETFRNLKYEVRNKNDLTREEIVELMRDVSKEDHSKRSSFVCVLLSHGEEGIIFGTNGPVDLKKITNFFRGDRCRSLTGKPKLFIIQACRGTELDCGIETDSGVDDDMACHKIPVEADFLYAYSTAPGYYSWRNSKDGSWFIQSLCAMLKQYADKLEFMHILTRVNRKVATEFESFSFDATFHAKKQIPCIVSMLTKELYFYSGGGSGGGGENLYFQSGGGPAGEASSIPNREGKPIPNPLLGLGSTRTGEFEIGTGFPFDPHYVEVLGERMHYVDVGPRDGTPVLFLHGNPTSSYVWRNIIPHVAPTHRCIAPDLIGMGKSDKPDLGYFFDDHVRFMDAFIEALGLEEVVLVIHDWGSALGFHWAKRNPERVKGIAFMEFIRPIPTWDEWPEFARETFQAFRTTDVGRKLIIDQNVFIEGTLPMGVVRPLTEVEMDHYREPFLNPVDREPLWRFPNELPIAGEPANIVALVEEYMDWLHQSPVPKLLFWGTPGVLIPPAEAARLAKSLPNCKAVDIGPGLNLLQEDNPDLIGSEIARWLSTLEISGHHHHHH<br>HaloTag/TVMV-CS/Casp3/TEV-CS/HaloTag/His$_6$-Tag | Amino acid sequence for Caspase-3 construct, which is N-/ and C-/terminally fused to a HaloTag via a long linker containing TVMV-CS and TEV-CS on the N-/ and C-/terminal, respectively. | This work |
| **pHM118**<br><br>**(split-TEV)** | MGSAWSHPQFEKGGGSGGGSGGSAWSHPQFEKSAGKSMSSMVSDTSCTFPSSDGIFWKHWIQTKDGQCGSPLVSTRDGFIVGIHSASNFTNTNN | Amino acid sequence for C-terminal Twin- | This work |

| | | | |
|---|---|---|---|
| | YFTSVPKNFMELLTNQEAQQWVSGWRLNADSVLWGGHKVFM<br>VGGGSGGGDEVDGGGSGGGSPEDELAANEEELQQNEQKLAQI<br>KQKLQAIKYGGGSGGDEVDGGSGGSPEDEIQQLEEEIAQLEQK<br>NAALKEKNQALKYGGSGGSGGSGGSGGSGGGESLFKGPRDYN<br>PISSTICHLTNESDGHTTSLYGIGFGPFIITNKHLFRRNNGTLLVQ<br>SLHGVFKVKNTTTLQQHLIDGRDMIIIRMPKDFPPFPQKLKFREP<br>QREERICLVTTNFQTGSGSGSGSGSGSNSYDSSSIKVLKGLDAV<br>RKRPGMYIGDTDDGTGLHHMVFEVVDNAIDEALAGHCKEIIVT<br>IHADNSVSVQDDGRGIPTGIHPEEGVSAAEVIMTVLHAGGKFDD<br>NSYKVSGGLHGVGVSVVNALSQKLELVIQREGKIHRQIYEHGV<br>PQAPLAVTGETEKTGTMVRFWPSLETFTNVTEFEYEILAKRLRE<br>LSFLNSGVSIRLRDKRDGKEDHFHYEG<br>Twin-StrepTag/cTEV/CCS/AP4/CCS/P3/nTEV/GyrB | StrepTag for purification, C-terminal split-TEV fused to AP4 coiled region via a linker containing CCS, followed by another CCS-containing linker fusing P3 coiled region to N-terminal of split-TEV. | |
| **pHM69** | MKIEEGKLVIWINGDKGYNGLAEVGKKFEKDTGIKVTVEHPDK<br>LEEKFPQVAATGDGPDIIFWAHDRFGGYAQSGLLAEITPDKAFQ<br>DKLYPFTWDAVRYNGKLIAYPIAVEALSLIYNKDLLPNPPKTWE<br>EIPALDKELKAKGKSALMFNLQEPYFTWPLIAADGGYAFKYEN<br>GKYDIKDVGVDNAGAKAGLTFLVDLIKNKHMNADTDYSIAEA<br>AFNKGETAMTINGPWAWSNIDTSKVNYGVTVLPTFKGQPSKPF<br>VGVLSAGINAASPNKELAKEFLENYLLTDEGLEAVNKDKPLGA<br>VALKSYEEELAKDPRIAATMENAQKGEIMPNIPQMSAFWYAVR<br>TAVINAASGRQTVDEALKDAQTNSSSNNNNNNNNGGGSGGGS<br>GGGSSKALLKGVRDFNPISACVCLLENSSDGHSERLFGIGFGPYI<br>IANQHLFRRNNGELTIKTMHGEFKVKNSTQLQMKPVEGRDIIVI<br>KMAKDFPPFPQKLKFRQPTIKDRVCMVSTNFQQKSVSSLVSESS<br>HIVHKEDTSFWQHWITTKDGQCGSPLVSIIDGNILGIHSLTHTTN<br>GSNYFVEFPEKFVATYLDAADGWCKNWKFNADKISWGSFTLV<br>EDAPEDHHHHHH<br>Maltose-binding-protein (MBP)/TVMV/His$_6$-Tag | MBP fused to TVMV protease (full version) and a His$_6$-Tag for purification | This work |
| **pRK793** | MKTEEGKLVIWINGDKGYNGLAEVGKKFEKDTGIKVTVEHPD<br>KLEEKFPQVAATGDGPDIIFWAHDRFGGYAQSGLLAEITPDKAF<br>QDKLYPFTWDAVRYNGKLIAYPIAVEALSLIYNKDLLPNPPKT<br>WEEIPALDKELKAKGKSALMFNLQEPYFTWPLIAADGGYAFKY<br>ENGKYDIKDVGVDNAGAKAGLTFLVDLIKNKHMNADTDYSIA<br>EAAFNKGETAMTINGPWAWSNIDTSKVNYGVTVLPTFKGQPSK<br>PFVGVLSAGINAASPNKELAKEFLENYLLTDEGLEAVNKDKPL<br>GAVALKSYEEELAKDPRIAATMENAQKGEIMPNIPQMSAFWYA<br>VRTAVINAASGRQTVDEALKDAQTNSSSNNNNNNNNNNLGIEG<br>RGENLYFQGHHHHHHHGESLFKGPRDYNPISSTICHLTNESDGH<br>TTSLYGIGFGPFIITNKHLFRRNNGTLLVQSLHGVFKVKNTTTLQ<br>QHLIDGRDMIIIRMPKDFPPFPQKLKFREPQREERICLVTTNFQT<br>KSMSSMVSDTSCTFPSSDGIFWKHWIQTKDGQCGSPLVSTRDG<br>FIVGIHSASNFTNTNNYFTSVPKNFMELLTNQEAQQWVSGWRL<br>NADSVLWGGHKVFMVKPEEPFQPVKEATQLMNRRRRR<br>MBP/ TEV-CS /His$_6$-Tag/TEV | MBP fused to TEV protease (full version) as a solubility tag. His$_6$-Tag is placed between MBP and TEV sequence to enable purification of TEV without MBP-Tag. | [2] |

**Table S2.** Protein production conditions of different constructs.

| Protein | Temperature (°C) | Incubation time (h) | Purification resin |
|---|---|---|---|
| Split-TEV | 30 | 4 | Strep-Tactin®XT |
| Split-TVMV | 30 | 3.5 | $Ni^{2+}$-NTA |
| Casp3 | 18 | 16 | $Ni^{2+}$-NTA |
| Displacer | 30 | 4 | $Ni^{2+}$-NTA |
| MBP-TVMV | 30 | 4 | $Ni^{2+}$-NTA |
| TEV | 30 | 4 | $Ni^{2+}$-NTA |

**Table S3.** Protein identification and relative quantification by LC-MS/MS.

| Protein | Band MW [kDa] | Peptides | iBAQ | iBAQ rank | rel. iBAQ [%] | Sequence Coverage [%] |
|---|---|---|---|---|---|---|
| pHM111 | 105 | 49 | 3.92 E8 | 1 | 37.8 | 56.2 |
| pHM118 | 70 | 58 | 5.07 E9 | 1 | 98.7 | 97.5 |
| pHM83 | 80 | 57 | 1.63 E9 | 1 | 94.5 | 71.6 |

#### References


[1] C. Geraths, M. Daoud-El Baba, G. Charpin-El Hamri, W. Weber, *J. Control. Release* **2013**, *171*, 57.

[2] R. B. Kapust, J. Tözsér, J. D. Fox, D. E. Anderson, S. Cherry, T. D. Copeland, D. S. Waugh, *Protein Eng. Des. Sel.* **2001**, *14*, 993.

[3] V. Landwehr, M. Milanov, L. Angebauer, J. Hong, G. Jüngert, A. Hiersemenzel, A. Siebler, F. Schmit, Y. Öztürk, S. Dannenmaier, F. Drepper, B. Warscheid, H. G. Koch, *Front. Mol. Biosci.* **2021**, *8*, 1.

### Supporting Information (Model Summary)

*Hasti Mohsesin, Hanna J. Wagner, Marcus Rosenblatt, Svenja Kemmer, Friedel Drepper, Pitter Huesgen, Jens Timmer* and *Wilfried Weber**

#### 1 Description of the mathematical model

##### 1.1 Mathematical model of the decoder

We established a mathematical model to characterize, predict and optimize the functionality of the designed biohybrid binary decoder. The model as summarized in Tables 1 and 2 consists of ordinary differential equations (ODEs) describing the dynamical behavior of the system's components TEV (module A), TVMV (module B) and Casp3 (module C). Each of these components appears in three different configurations with the compound being bound (*bd*) to the magnetic beads, released and active (*on*) as well as released and inactive (*off*). Via the reactions $v_1$ and $v_8$ initiated by novobiocin or EDTA, TEV and TVMV are released from the magnetic bead and present in the active form, respectively. From the active form, they can be deactivated (reactions $v_2$ and $v_9$) and TEV can also be reactivated ($v_4$). The deactivation of TEV is enhanced by the displacer which is formulated as a power law in reaction $v_3$. When active Casp3 is present, TEV and TVMV can go over into an intermediate configuration *cut* (reactions $v_6$, $v_7$, $v_{10}$ and $v_{11}$), again enhanced by the displacer for the case of TEV. From the *cut* state, the compounds can no longer be activated but relax into the corresponding *cut_off* state. Note that the *cut* state still contributes to the measured enzyme activity as shown by the observation function in Table 3, whereas the *cut_off* state does not contribute to the activity. The release of Casp3 by TEV and TVMV was modeled as a two-step process to allow for both orders of either TEV being present first followed by TVMV or the opposite way. All reactions of the ODE system were derived from the law of mass action including Michaelis-Menten kinetics. For the displacer, at first a Hill-type kinetic was assumed which resulted in a power law after model reduction [5].

##### 1.2 Parameter estimation and identifiability analysis

Since it is difficult to measure the kinetic rates of the established system directly, these were determined by means of a mathematical modeling approach using the available measurement data and maximum likelihood estimation. The ODE system is written as $\frac{d}{dt}x(t) = f(x(t), p, u(t)$, with internal states $x$ and unknown dynamic parameters $p$, e.g. biochemical rates. Initial conditions of the states are also parameters, termed $x_0 = x(t = 0)$. The full list of equations is written in Table 1 and Table 2 shows how the individual reactions add up to the time derivatives of the states, i.e. how the states vary over time. Experimental conditions such as adding a compound to the system at a particular time point are modeled via an input function $u(t)$ which is part of the ODE system. To link the model's time courses to experimental data, we use an observation function $y = g(x, p)$ that includes scaling and offset parameters (see Table 3). The measurement error was assumed to be Gaussian normally distributed and was described by an absolute error model including error parameters (see Table 3).
Model parameters were determined via maximum likelihood estimation from experimental data using the modeling environment `dMod` [1] in `R`. Following state-of-the-art methodology [2], a deterministic multi-start optimization was performed starting from 1000 randomly chosen positions in parameter space. A number of 62 optimization runs converged to the global optimum as illustrated by a waterfall plot in Figure 1. Parameter values of the best resulting fit are summarized in Table 4. To determine confidence intervals of all parameters of the system, we computed for each parameter the corresponding profile likelihood [3] as shown in Figures 2 and 3. In order to simplify parameter estimation and obtain meaningful predictions, the ODE model undervent several steps of model refinement including identifiability analysis and model reduction [4].

#### 1.3 Incorporation different experimental conditions and datasets

As a particular focus, the medium change and incubation time of the measurement sample was explicitely modeled. In the modeling framework, a condition-specific event time point was included at which several model parameters were allowed to take different values (compare Table 4) than before to account for the new experimental conditions of incubation at a temperature of $4°C$. Figure 4 exemplarily shows how the model trajectories change due to this medium change and how this influenced the model calibration. For the calibration of the ODE model, two datasets were used that differed in their corresponding experimental settings. Since these datasets had not been measured on a compatible scale, different scaling, offset and error parameters were for the two data sets (compare Table 3).

#### 1.4 Model prediction and validation

To test the predictive power of the established ODE model, different displacer doses and incubation times were simulated (compare main text Figure 3d). Based on this result, an incubation time of $t = 150\,\mathrm{min}$ and a displacer concentration of $0.087\frac{mg}{ml}$ was chosen for a subsequent time course experiment. The established ODE model was applied to the newly obtained experimental data, whereby only new scaling and offset parameters were estimted, but dynamic parameters were taken from the best fit to the original data set. Again, uncertainties of the experimental data was estimated using an error model. Results of the model validation are shown in main text Figures 4e and S5.

#### References

[1] D. Kaschek, W. Mader W, M. Fehling-Kaschek, M. Rosenblatt, J. Timmer. Dynamic modeling, parameter estimation, and uncertainty analysis in R. *Journal of Statistical Software* **2019** 88(10), 1–32
[2] A. Raue, M. Schilling, J. Bachmann, A. Matteson, M. Schelker, D. Kaschek, S. Hug, C. Kreutz, B.D. Harms, F.J. Theis, U. Klingmüller, J. Timmer. Lessons learned from Quantitative Dynamical Modeling in Systems Biology. *PLOS ONE* **2013** 8(12)
[3] A. Raue, C. Kreutz, T. Maiwald, J. Bachmann, M. Schilling, U. Klingmüller, J. Timmer *Bioinformatics* **2009** 25(15)
[4] F.-G. Wieland, A. Hauber, M. Rosenblatt, C. Tönsing, J. Timmer. On Structural and Practical Identifiability. *Current Opinion in Systems Biology* **2021**
[5] T. Maiwald, H. Hass, B. Steiert, J. Vanlier, R. Engesser, A. Raue, F. Kipkeew, H.H. Bock, D. Kaschek, c. Kreutz, J. Timmer. Driving the model to its limit: Profile Likelihood based Model Reduction. *PLOS ONE* **2016**

Table 1: Reactions of the ODE model are based on the law of mass action, Michaelis-Menten and Hill-kinetic terms where necessary. The table shows the final model after model reduction [5].

| Reaction | Educt | Product | Rate | Description |
|---|---|---|---|---|
| $v_1$ | $TEV_{bd}$ | $TEV_{on}$ | novo · k_rel_TEV · $TEV_{bd}$/(Km_novo + $TEV_{bd}$) | TEV release by novobiocin |
| $v_2$ | $TEV_{on}$ | $TEV_{off}$ | k_inact_TEV_basal · $TEV_{on}$ | TEV basal inactivation |
| $v_3$ | $TEV_{on}$ | $TEV_{off}$ | k_disp_inact_TEV_basal · $TEV_{on}$ · $Disp^{n\_inact\_TEV\_basal}$ | TEV basal inactivation enhanced by Displacer |
| $v_4$ | $TEV_{off}$ | $TEV_{on}$ | k_react_TEV · $TEV_{off}$ | TEV reactivation |
| $v_5$ | $TEV_{on}$ | $TEV_{cut}$ | k_cut_TEV_by_Casp3 · $Casp3_{on}$ · $TEV_{on}$· 1/(Km_Casp3TEV + $TEV_{on}$)/(Km_TEVCasp3 + Casp3_on) | TEV cutting by Casp3 |
| $v_6$ | $TEV_{cut}$ | $TEV_{cut\_off}$ | k_inact_TEV · $TEV_{cut}$ | TEV inactivation |
| $v_7$ | $TEV_{cut}$ | $TEV_{cut\_off}$ | $10^6$ · k_disp_inact_TEV · $TEV_{cut}$ · $Disp^{n\_inact\_TEV}$ | TEV inactivation enhanced by Displacer |
| $v_8$ | $TVMV_{bd}$ | $TVMV_{on}$ | EDTA · $k_{rel}$TVMV · $TVMV_{bd}$/ (KmEDTA + $TVMV_{bd}$) | TVMV release by EDTA |
| $v_9$ | $TVMV_{on}$ | $TVMV_{off}$ | k_inact_TVMV_basal · $TVMV_{on}$ | TVMV basal inactivation |
| $v_{10}$ | $TVMV_{on}$ | $TVMV_{cut}$ | k_cut_TVMV_byCasp3 · $Casp3_{on}$ · $TVMV_{on}$· 1/(KmCasp3TVMV + $TVMV_{on}$) | TVMV cutting by Casp3 |
| $v_{11}$ | $TVMV_{on}$ | $TVMV_{cut}$ | k_cut_TVMV_byCasp3bdTVMVCS · $Casp3_{bd}$TVMVCS· $TVMV_{on}$/(KmCasp3TVMV + $TVMV_{on}$) | TVMV cutting by $Casp3_{bd}TVMV_{CS}$ |
| $v_{12}$ | $TVMV_{cut}$ | $TVMV_{cut\_off}$ | k_inact_TVMV · $TVMV_{cut}$ | TVMV inactivation |
| $v_{13}$ | $TVMV_{off}$ | $TVMV_{on}$ | k_react_TVMV · $TVMV_{off}$ | TVMV reactivation |
| $v_{14}$ | $Casp3_{bd}$ | $Casp3_{bd}$TVMVCS | k_cut_Casp3_TEV · ($TEV_{on}$ + $TEV_{cut}$) · $Casp3_{bd}$ | $Casp3_{bd}$ cut by TEV |
| $v_{15}$ | $Casp3_{bd}$ | $Casp3_{bd}$TEVCS | k_cut_Casp3_TVMV · ($TVMV_{on}$ + $TVMV_{cut}$)· $Casp3_{bd}$/(KmTVMV + $Casp3_{bd}$) | $Casp3_{bd}$ cut by TVMV |
| $v_{16}$ | $Casp3_{bd}$TEVCS | $Casp3_{on}$ | k_cut_Casp3_TEV · ($TEV_{on}$ + $TEV_{cut}$) · $Casp3_{bd}$TEVCS | $Casp3_{bd}$TEVCS cut by TEV |
| $v_{17}$ | $Casp3_{bd}$TVMVCS | $Casp3_{on}$ | k_cut_Casp3_TVMV · ($TVMV_{on}$ + $TVMV_{cut}$)· $Casp3_{bd}$TVMVCS/(KmTVMV + $Casp3_{bd}$TVMVCS) | $Casp3_{bd}$TVMVCS cut by TVMV |
| $v_{18}$ | $Casp3_{bd}$TEVCS | $Casp3_{on}$ | k_rel_Casp3bdTEVCS_leaky · $Casp3_{bd}$TEVCS | $Casp3_{bd}$TEVCS leaky release |
| $v_{19}$ | $Casp3_{bd}$TVMVCS | $Casp3_{on}$ | k_rel_Casp3bdTVMVCS_leaky · $Casp3_{bd}$TVMVCS | $Casp3_{bd}$TVMVCS leaky release |
| $v_{20}$ | $Casp3_{on}$ | $Casp3_{off}$ | k_inact_Casp3 · $Casp3_{on}$ | Casp3 inactivation |

Table 2: Ordinary differential equations of the decoder model.

| Time derivative | | ODE's right hand side | Initial value at $t = 0$ |
|---|---|---|---|
| $\frac{d}{dt}\text{TEV}_{\text{bd}}$ | $=$ | $-v_1$ | TEV_bd_0 |
| $\frac{d}{dt}\text{TEV}_{\text{on}}$ | $=$ | $v_1 - v_2 - v_3 + v_4 - v_5$ | 0 |
| $\frac{d}{dt}\text{TEV}_{\text{off}}$ | $=$ | $v_2 + v_3 - v_4$ | 0 |
| $\frac{d}{dt}\text{TEV}_{\text{cut}}$ | $=$ | $v_5 - v_6 - v_7$ | 0 |
| $\frac{d}{dt}\text{TEV}_{\text{cut\_off}}$ | $=$ | $v_6 + v_7$ | 0 |
| $\frac{d}{dt}\text{TVMV}_{\text{bd}}$ | $=$ | $-v_8$ | TVMV_bd_0 |
| $\frac{d}{dt}\text{TVMV}_{\text{on}}$ | $=$ | $v_8 - v_9 - v_{10} - v_{11}$ | 0 |
| $\frac{d}{dt}\text{TVMV}_{\text{off}}$ | $=$ | $v_9 - v_{13}$ | 0 |
| $\frac{d}{dt}\text{TVMV}_{\text{cut}}$ | $=$ | $v_{10} + v_{11} - v_{12}$ | 0 |
| $\frac{d}{dt}\text{TVMV}_{\text{cut\_off}}$ | $=$ | $v_{12}$ | 0 |
| $\frac{d}{dt}\text{Casp3}_{\text{bd}}$ | $=$ | $-v_{14} - v_{15}$ | Casp3_bd_0 |
| $\frac{d}{dt}\text{Casp3}_{\text{bd}}\text{TEVCS}$ | $=$ | $v_{15} - v_{16} - v_{18}$ | 0 |
| $\frac{d}{dt}\text{Casp3}_{\text{bd}}\text{TVMVCS}$ | $=$ | $v_{14} - v_{17} - v_{19}$ | 0 |
| $\frac{d}{dt}\text{Casp3}_{\text{on}}$ | $=$ | $v_{16} + v_{17} + v_{18} + v_{19} - v_{20}$ | 0 |
| $\frac{d}{dt}\text{Casp3}_{\text{off}}$ | $=$ | $v_{20}$ | 0 |

Table 3: Observation function of the ODE model. For each measured compound, a scaling factor and an offset is assumed. For TEV and TVMV, the *cut* compound contributes with a different scaling factor.

| Observable | Computation from model states | Error parameter |
|---|---|---|
| $\text{TEV}_{\text{obs}}$ | $\text{TEV}_{\text{on}} \cdot \text{sc\_TEV} + \text{TEV}_{\text{cut}} \cdot \text{sc\_TEV} \cdot \text{sc\_TEV\_cut\_factor} + \text{off\_TEV}$ | sd_TEV |
| $\text{TVMV}_{\text{obs}}$ | $\text{TVMV}_{\text{on}} \cdot \text{sc\_TVMV} + \text{TVMV}_{\text{cut}} \cdot \text{sc\_TVMV} \cdot \text{sc\_TVMV\_cut\_factor} + \text{off\_TVMV}$ | sd_TVMV |
| $\text{Casp3}_{\text{obs}}$ | $\text{Casp3}_{\text{on}} \cdot \text{sc\_Casp3} + \text{off\_Casp3}$ | sd_Casp3 |

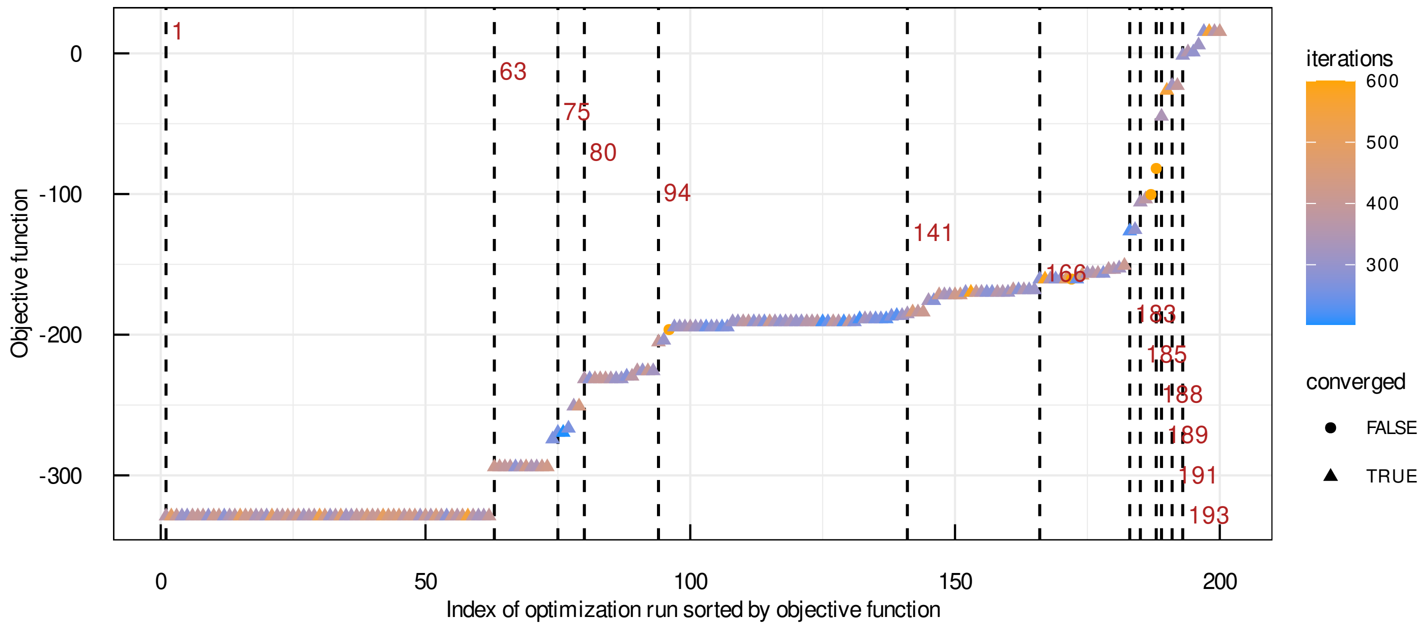


Figure 1: Optimization runs sorted by the objective function displayed as a waterfall plot. The best 200 out of 1000 optimization runs are shown. Steps correspond to local optima. The global optimum was found in 62 of the 1000 optimzation runs.

Table 4: Parameter values of the best fit. For each parameter, lower and upper bounds of the 68%-confidence intervals are shown as computed from the profile likelihood.

| Number | Name of parameter | Estimated | Scale | bestfit value | lower bound | upper bound |
|---|---|---|---|---|---|---|
| 1 | k_cut_Casp3_TEV | True | log10 | -3.149 | -3.523 | -2.954 |
| 2 | k_cut_Casp3_TVMV | True | log10 | 1.402 | 0.877 | 3.751 |
| 3 | k_cut_TEV_byCasp3 | True | log10 | -3.062 | -3.246 | -2.838 |
| 4 | k_cut_TVMV_byCasp3TVMVCS | True | log10 | -2.253 | -2.922 | -1.686 |
| 5 | k_disp_k_inact_TEV | True | log10 | 3.060 | 2.929 | 3.267 |
| 6 | k_disp_k_inact_TEV_basal | True | log10 | -0.0657 | -0.246 | 0.443 |
| 7 | k_inact_Casp3 | True | log10 | -1.875 | -2.250 | -1.609 |
| 8 | k_inact_TEV_basal | True | log10 | -2.371 | -2.541 | -2.235 |
| 9 | k_inact_TVMV | True | log10 | -2.045 | -2.540 | -1.706 |
| 10 | k_inact_TVMV_basal | True | log10 | -0.264 | -0.592 | 0.0448 |
| 11 | k_react_TVMV | True | log10 | -2.324 | -2.380 | -2.2784 |
| 12 | k_rel_Casp3TEVCS_leaky | True | log10 | -2.442 | -2.682 | -2.069 |
| 13 | k_rel_Casp3TVMVCS_leaky | True | log10 | -1.881 | -2.150 | -1.558 |
| 14 | k_rel_TEV | True | log10 | -2.379 | -2.564 | -2.175 |
| 15 | k_rel_TVMV | True | log10 | -4.821 | -4.857 | -4.778 |
| 16 | KmCasp3_TEV | True | log10 | -3.053 | -Inf | -1.921 |
| 17 | KmCasp3_TVMV | True | log10 | -3.607 | -4.452 | -2.849 |
| 18 | KmEDTA | True | log10 | -3.699 | -3.926 | -3.492 |
| 19 | KmNovo | True | log10 | -1.107 | -1.568 | -0.673 |
| 20 | KmTEV_Casp3 | True | log10 | -2.146 | -2.649 | -1.752 |
| 21 | KmTVMV | True | log10 | -0.461 | -1.321 | 2.676 |
| 22 | NM_k_cut_TVMV_byCasp3 | True | log10 | -1.531 | -1.952 | -1.307 |
| 23 | NM_k_inact_Casp3 | True | log10 | -2.454 | -2.635 | -2.326 |
| 24 | NM_k_inact_TEV | True | log10 | -1.560 | -1.660 | -1.448 |
| 25 | NM_k_inact_TEV_basal | True | log10 | -1.153 | -1.293 | -0.669 |
| 26 | NM_k_inact_TVMV_basal | True | log10 | -1.180 | -1.390 | -0.9355 |
| 27 | NM_k_react_TEV | True | log10 | -1.259 | -1.415 | -0.756 |
| 28 | off_Casp3_ds1 | True | log10 | -0.00288 | -0.060 | 0.016 |
| 29 | off_Casp3_ds2 | True | log10 | -0.314 | -0.353 | -0.276 |
| 30 | off_TEV_ds2 | True | log10 | -0.533 | -0.593 | -0.480 |
| 31 | off_TVMV_ds1 | True | log10 | -0.241 | -1.089 | -0.151 |
| 32 | off_TVMV_ds2 | True | log10 | 0.083 | 0.064 | 0.102 |
| 33 | sc_Casp3_ds2 | True | log10 | -0.829 | -1.059 | -0.446 |
| 34 | sc_TEV_cut_factor | True | log10 | 0.319 | 0.264 | 0.380 |
| 35 | sc_TEV_ds1 | True | log10 | -0.941 | -1.004 | -0.894 |
| 36 | sc_TEV_ds2 | True | log10 | -1.371 | -1.507 | -1.225 |
| 37 | sc_TVMV_cut_factor | True | log10 | -1.754 | -1.993 | -1.563 |
| 38 | sc_TVMV_ds1 | True | log10 | -0.438 | -0.730 | 0.134 |
| 39 | sc_TVMV_ds2 | True | log10 | 2.731 | 2.420 | 3.045 |
| 40 | sd_Casp3_ds1 | True | log10 | -1.197 | -1.353 | -0.999 |
| 41 | sd_Casp3_ds2 | True | log10 | -0.751 | -0.804 | -0.695 |
| 42 | sd_TEV_ds1 | True | log10 | -0.917 | -1.001 | -0.821 |
| 43 | sd_TEV_ds2 | True | log10 | -0.805 | -0.857 | -0.748 |
| 44 | sd_TVMV_ds1 | True | log10 | -0.710 | -0.802 | -0.612 |
| 45 | sd_TVMV_ds2 | True | log10 | -0.670 | -0.722 | -0.610 |
| 46 | TEV_bd_0 | True | log10 | -0.617 | -0.766 | -0.481 |
| 47 | TVMV_bd_0 | False | lin | 0.001 | NA | NA |
| 48 | Casp3_bd_0 | False | lin | 1 | NA | NA |
| 49 | k_cut_TVMV_byCasp3 | False | lin | 0 | NA | NA |
| 50 | k_inactivate_TEV | False | lin | 0 | NA | NA |
| 51 | k_reactivate_TEV | False | lin | 0 | NA | NA |
| 52 | n_k_inactivate_TEV | False | lin | 1.564 | NA | NA |
| 53 | n_k_inactivate_TEV_basal | False | lin | 8 | NA | NA |
| 54 | off_TEV_ds1 | False | lin | 0 | NA | NA |

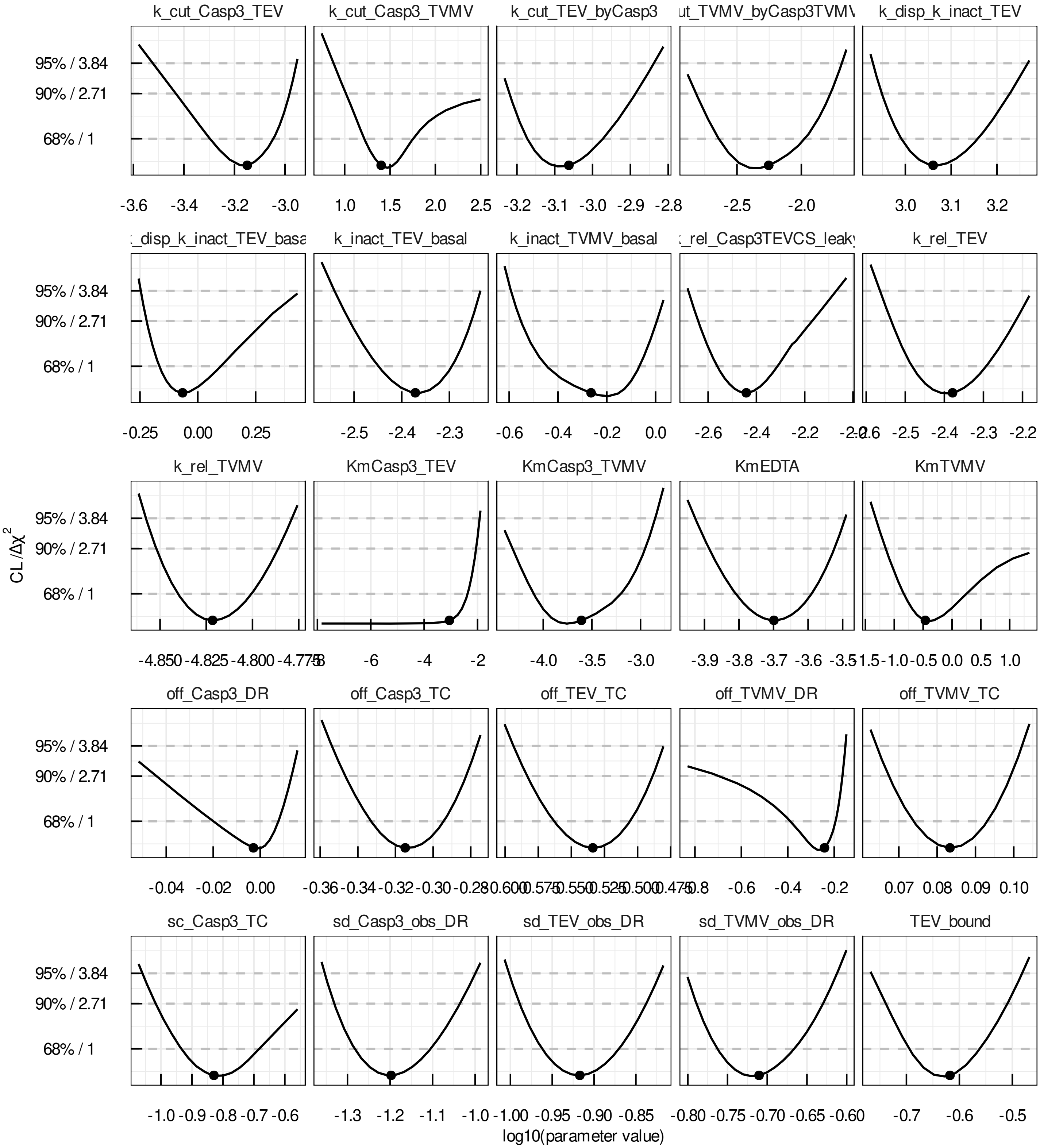


Figure 2: Parameter profile likelihood [3] computed for parameters 1-25 of the best parameter set. All parameters are identifiable except for KmCasp3_TEV whose parameter value is in agreement with minus infinity. The parameter was not reduced from the model, to conserve the mass-action based formulation of the model.

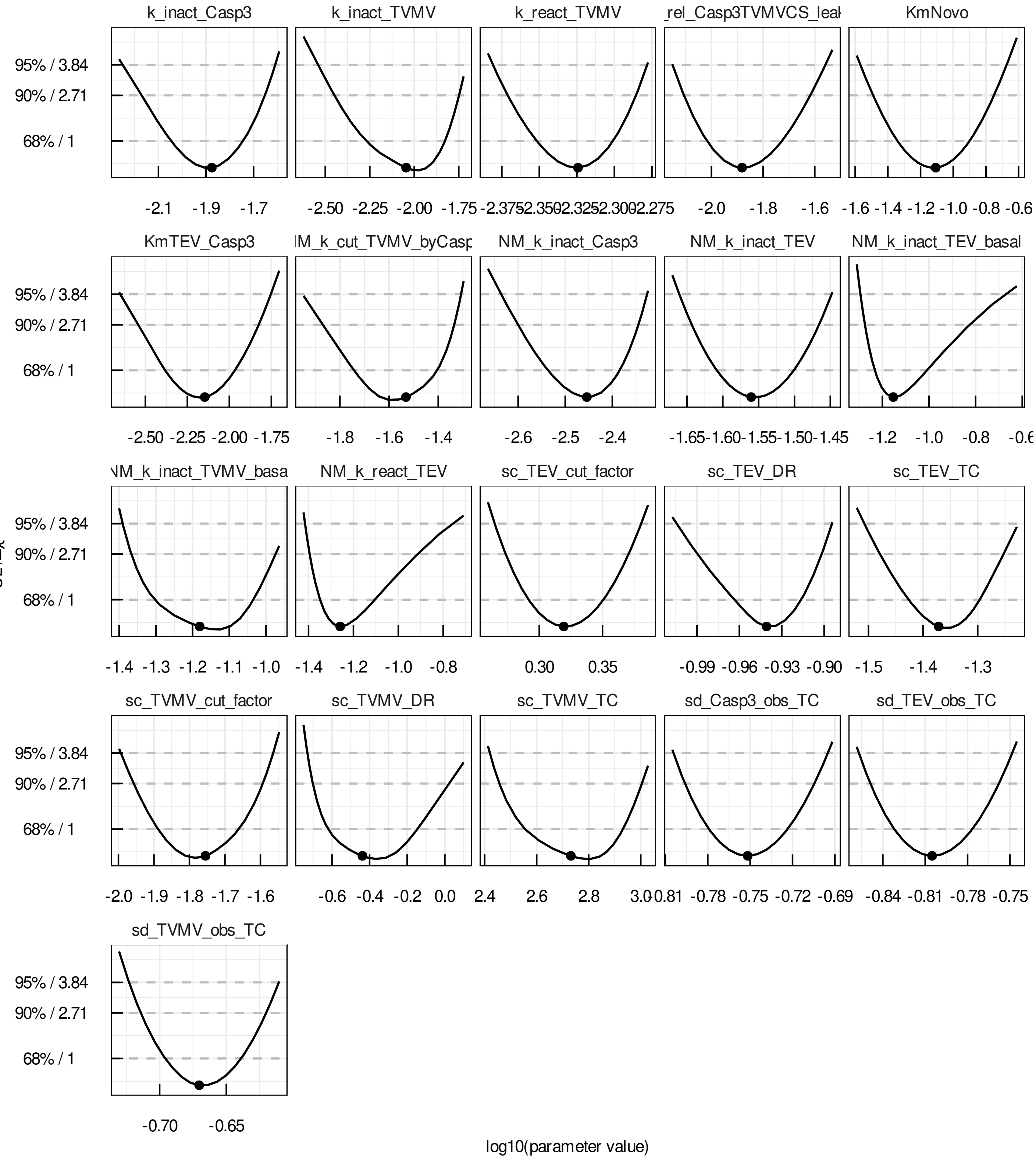


Figure 3: Parameter profile likelihood [3] computed for parameters 26-46 of the best parameter set. All parameters are identifiable.

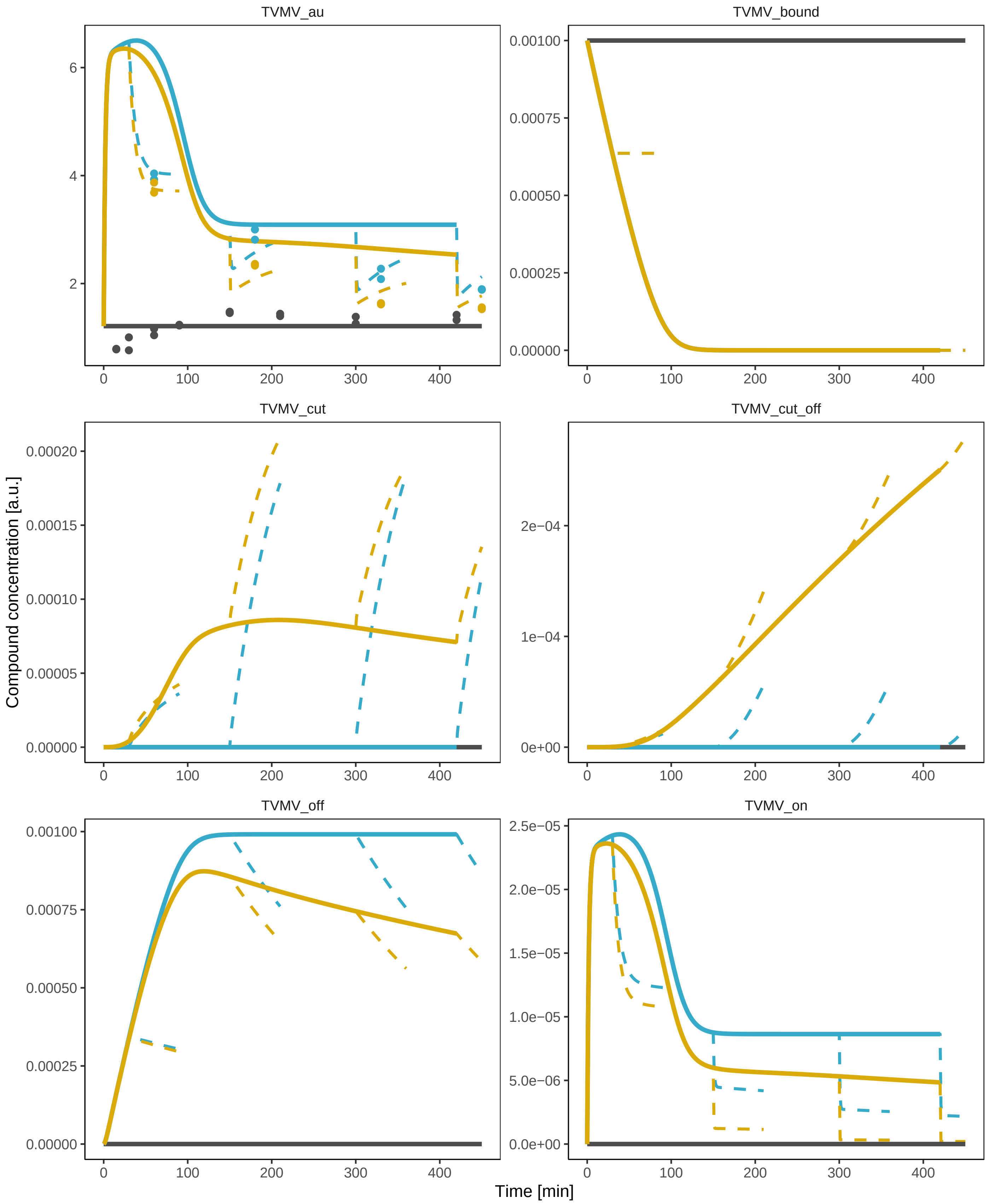


Figure 4: Exemplary model trajectories for TVMV compounds influenced by medium change. The control condition (black) is shown together with EDTA (yellow) and novobiocin+EDTA (blue) conditions. Measurement data (dots) and model trajectories of first medium (solid lines) and second medium (dashed line) are shown. For simplification, only four different measurement time points after incubation are shown. The modeled medium change is visible by a sharp change in the model's time courses.
